# Distinct decision processes for 3D and motion stimuli in both humans and monkeys revealed by computational modelling

**DOI:** 10.1101/2024.11.25.625189

**Authors:** Revan Rangotis, Sabina Nowakowska, Peter Dayan, Andrew J. Parker, Ehsan Kakaei, Abibat Akande, Kristine Krug

**Affiliations:** Institute of Biology, Otto-von-Guericke-University Magdeburg, Magdeburg, Germany; University Medical Centre Göttingen, Institute for Auditory Neuroscience, Göttingen, Germany; Max Planck Institute for Biological Cybernetics & University of Tübingen, Tübingen, Germany; Department of Physiology Anatomy and Genetics, University of Oxford, Oxford, United Kingdom; Leibniz Institute for Neurobiology, Magdeburg, Germany

## Abstract

Decision-making models distil principles of information processing that underlie a range of cognitive functions. For the cases of motion detection in random-dot kinematograms and decisions about the rotation of 3D structure-from-motion cylinders, contributing neural processes have been localized to specific circuits in extrastriate area V5/MT on the basis of both causal and correlative evidence, suggesting a common decision path. Here, we arranged for humans and rhesus monkeys to make perceptual choices about these two stimulus types, indicating their decision with either hand or eye movements. In both species, the parameter distributions of a hierarchical Drift Diffusion Model (DDM) revealed systematic differences for stimulus type but not the different modes of response. States in a hidden Markov model also indicated distinct decision strategies, again for the two stimulus types but not for response mode. Although physiological evidence points to area V5/MT as a vehicle for relevant perceptual signals for both stimulus types, computational modelling of the present results reveal distinct decision processes, therefore predicting that different neural processes underlie judgements about motion and 3D-depth, consistently for humans and monkeys.

## Introduction

The framework of sequential sampling has provided a powerful collection of models of sensory, affective and memory-based decision-making. The principal goal of these models is to use behavioural data, in particular response accuracy and reaction time (RT), to make inferences about aspects of the neural computations and mechanisms that underlie critical cognitive processes (Ratcliff & McKoon, 2008). Substrates linked to model parameters have been proposed in humans (e.g. Heekeren et al., 2004) and macaque monkeys (e.g. Roitman & Shadlen, 2002).

There is a variety of popular models (Brown & Heathcote, 2008; Krajbich et al., 2010; Usher & McClelland, 2001), many of which are based on a noisy evidence accumulation process, which terminates once a threshold, or “bound”, is reached (Forstmann et al., 2016). In particular, the drift diffusion model (DDM) has become widely used for two-alternative forced choice tasks (2AFC) (Bogacz et al., 2006; Ratcliff, 1978, Ratcliff et al., 2016). While DDMs have been used extensively in different species (e.g., (Fan et al., 2018)), few studies have directly compared model performance across decision tasks or across species on the same task (but see (Voss et al., 2004)). Such comparisons are essential if we are to generalise our understanding of cognitive processes within single brains and between species, especially from animal models to humans (Nestler & Hyman, 2010). The models propose a pipeline from sensory input, through evidence inference and integration, to action selection and execution. There is a need to understand which neural elements are shared between tasks and which elements or processes are specific to particular tasks.

An assumption that is embedded in the use of the DDM is that the detailed strategy employed by a subject is fixed across an experimental session. There is ample empirical evidence that this is not true, e.g., (Braun et al., 2018; Purcell & Kiani, 2016; Roy et al., 2021), suggesting problems for parameter inference when using the DDM alone. Models have been developed in which the current strategy is characterized as a latent state that determines response properties. Strategies may then be altered spontaneously (Ashwood et al., 2022) or in reaction to experimental circumstances (Mohammadi et al., 2024). In one prominent example (Ashwood et al., 2022), the responses made during the dominance of any single strategy are described by a psychometric function derived from the parameters of a generalized linear model (GLM), and the switching process is described by a hidden Markov model (HMM). Here, we use a combination of the DDM and the GLM-HMM to elucidate the cognitive processes and strategies that govern two perceptual decision tasks in monkeys and humans.

Noisy random dot kinematograms (RDK) have been extensively used to investigate mechanisms underlying motion perception (Britten et al., 1992), perceptual decision-making (Roitman & Shadlen, 2002), and perceptual learning (Zohary, Celebrini, et al., 1994). Similarly, 3D structure-from-motion (SfM) cylinders have been employed to examine the eponymous processes. A SfM cylinder is a bistable stimulus with competing, mutually-exclusive rotational percepts, which flip stochastically and can be disambiguated by the addition of 3D depth in the form of binocular disparity (Parker & Krug, 2003).

Examining the processing of these two stimulus classes should offer useful clues to the structure of the decision-making pipeline, since perceptual signals for motion and binocular disparity, as well as the combination of these two visual cues, have been identified in single neurons in the visual cortical area V5/MT of the dorsal stream in the primate (Born & Bradley, 2005; Bradley et al., 1998; Dodd et al., 2001; Krug, 2004). Causal evidence for the role of V5/MT in both tasks is provided by focal micro-stimulation, which reliably drives perceptual decisions for both RDK (Salzman et al., 1990) and SfM cylinders (Krug et al., 2013). Also for both the RDK (Britten et al., 1996) and the SfM cylinder (Dodd et al., 2001), there is correlative evidence of trial-by-trial neural activity predictive of future behavioural decisions. This predictive relationship is summarized by choice probability (CP), a measure of how reliably behavioural decisions can be predicted from neural activity. CP is higher in V5/MT for decisions about the SfM cylinder (Dodd et al., 2001) than the RDK (Britten et al., 1996); this higher CP is also associated with higher interneuronal correlations (Bair et al., 2001; Wasmuht et al., 2019; Zohary, Shadlen, et al., 1994). These differences in neuronal signalling are suggestive of distinct neuronal mechanisms existing within the same cortical area.

While the neural substrate for perceptual decisions about both motion and depth has been linked to V5/MT, neurophysiological results also indicate distinct circuitry for evidence accumulation depending on how the outcome of the perceptual decision is reported (Gold & Shadlen, 2007). Several candidate regions have been investigated, predominantly those involved in the selection and preparation of eye movements, including the superior colliculus (Horwitz & Newsome, 2001), dlPFC (Kim & Shadlen, 1999), LIP (Shadlen & Newsome, 2001), and FEF (Gold & Shadlen, 2000). In parietal cortex, lateral intraparietal area (LIP) shows signals related to evidence accumulation for both hand and eye responses, but the medial intraparietal area (MIP) shows more pronounced rising activity when hand responses are performed (de Lafuente et al., 2015). The processing in different cortical areas could indicate differences in decision processes.

Here, we investigate the perceptual decision processes underpinned by incoming sensory evidence for the cases of pure motion direction (RDK) or combinations of binocular disparity and motion (SfM). For each stimulus type, we modelled decisions that were reported either by eye movements or by hand responses **(Figure 1)**. We employed DDMs and GLM-HMMs to model the underlying decision processes and strategies for behavioural data from both monkeys and humans. We address two questions. Firstly, are there different perceptual decision processes depending on the visual cues to be judged and how the response is delivered, by an eye movement or a hand movement? Secondly, do the analytical models yield comparable results for humans and monkeys carrying out the same decision task?

**Figure 1.**
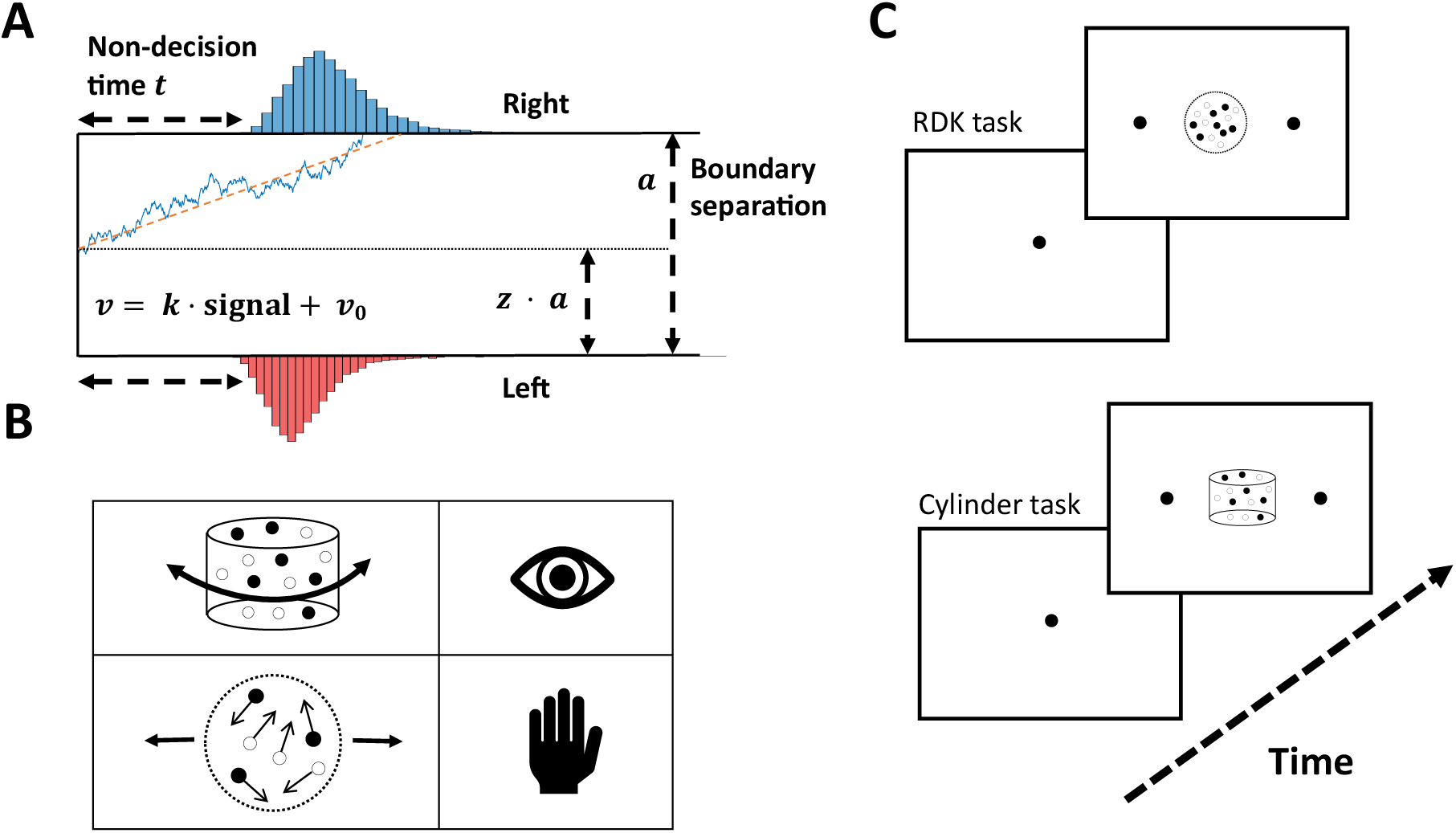
Task design and theoretical framework. **A**. Illustration of the trajectory of the decision process within the DDM framework with the key parameters and the distributions of reaction times for each decision. Non-decision time *t* is the mean minimum time for a participant to respond, related to stimulus encoding and motor processes. The evidence accumulation process, parametrised by the drift rate *v*, describing its slope, is given by the equation *v* = *k* ⋅ *signal* + *v*_0_ where “signal” is the normalised strength of the stimulus information, such as percentage coherence, and *v*_0_ is the intercept of the drift rate. *a* is the separation of the two decision boundaries, defining how much evidence is required before a decision is made. If *z* = 0.5, the decision boundaries are symmetrical around the starting point. The blue line represents evidence accumulation on one trial, the dashed orange line the average drift rate for one level of signed signal strength. **B**. We investigate the decision processes and behavioural strategies in perceptual judgements about the direction of motion in a random dot kinematogram (RDK) and about the rotation direction of a 3D structure-from-motion (SfM) cylinder in a reaction time (RT) task. In different experimental sessions, human and monkey participants reported their decisions on one of the two stimuli and with either a saccadic eye movement or a hand movement. We varied the stimulus information, i.e., motion coherence in the RDK and binocular disparity in the cylinder, across trials. **C**. This schematic illustrates the task design for eye movement responses. Upon successful fixation, an RDK or a 3D SfM cylinder stimulus was presented together with two choice targets, placed to the left and right of the stimulus. For hand movement responses, RDKs for the monkeys were also presented with two choice targets on a touchscreen, but for humans there were no choice targets on the screen as button presses were used to report instead.

## Results

20 humans and 2 rhesus monkeys (m133 and m134) made perceptual decisions about the direction of motion of an RDK and about the rotation direction of a 3D SfM cylinder. Subjects reported their decision with either an eye or a hand movement. The human participants carried out all four combinations of stimulus and response in a counterbalanced fashion. The monkeys performed the RDK task with hand responses and the cylinder task with saccade responses.

### Perceptual performance broadly comparable between humans and monkeys

We fitted behavioural responses from the RDK and cylinder tasks with cumulative Gaussian functions to determine the threshold and bias (**Figure 2A**). For the cylinder task, human participants performed similarly whether reporting by hand (mean threshold = 0.0086°, standard deviation (STD) = ±0.004°) or by eye (mean = 0.0084°, STD = ±0.005°) (n = 20, paired t-test, p = 0.7). This was also true for the RDK task (mean = 11.2%, STD = ±3.9% vs. mean = 13.3%, STD = ±5.0%) (n = 20, paired t-test, p = 0.1). Like humans, both monkeys had smooth psychophysical functions on both tasks. For the cylinder task, both monkeys had significantly higher thresholds than most humans (threshold m134 = 0.020°, m133 = 0.026°: single sample t-test, p = 0.0378 and p = 0.003, using methods described in (Crawford & Howell, 1998)). On the RDK task, monkey m134 showed similar accuracy to that of the humans (threshold = 12.3%, single sample t-test, p = 0.785), but m133’s threshold was considerably higher (threshold = 24.7%, single sample t-test, p = 0.003). However, m133’s threshold was still close to that of some human participants (**Figure 2A**).

**Figure 2.**
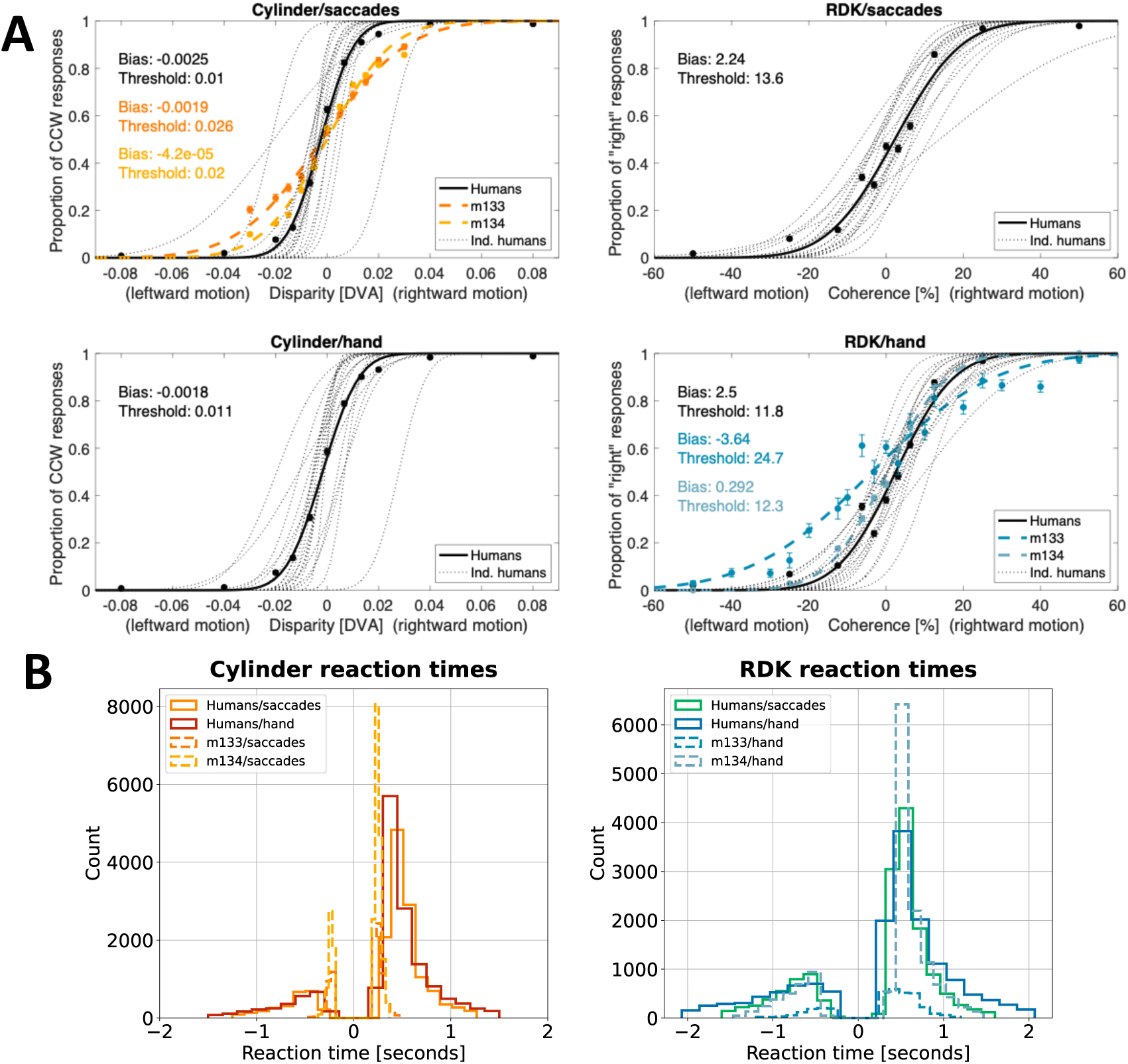
Psychophysical performance of humans and monkeys. **A**. We fitted cumulative Gaussian functions to the psychophysical response data separately for individual humans and for individual monkeys (m133, m134) for each perceptual decision task (3D SfM cylinder rotation; RDK direction of motion). For the cylinder, the binocular disparity units are degrees visual angle (DVA), while the RDK coherence is the percentage of the dots moving coherently in a specific direction. Overall, the performance of the humans (black) and monkeys (colored) is broadly comparable, but the monkeys’ thresholds were at the high end in comparison with the human sample. Individual data points correspond to proportions of choices pooled across sessions (monkeys) or across participants (humans) and the errors bars depict the SEM. **B**. Reaction time (RT) distributions for humans and monkeys. Negative reaction times represent incorrect responses. Solid lines show the pooled human participants (20), whereas the dashed lines show distribution for each of the monkeys pooled across all of their sessions. Again, the results are broadly comparable, but the monkeys made faster decisions, especially when they reported the rotation direction of the 3D SfM cylinder with an eye movement. For full RTs by stimulus strength see **Supplementary Figures 2-1** and for outlier removal **Supplementary Figures 2-2** and **2-3**.

Looking at the reaction time (RT) distributions (**Figure 2B**), while human and monkey participants carried out the perceptual tasks well, the monkeys tended generally to be faster but less accurate. A striking difference was that both monkeys were much faster responding with eye movements on the cylinder task than the human participants (mean RT = 0.252s, STD = ± 0.038s vs mean RT = 0.551s, STD = ± 0.234s; independent t-test, *p* < 0.0001), as previously observed for other saccade tasks (Middlebrooks & Schall, 2014). The monkeys, in particular, showed even faster error trials for the cylinder task (**Supplementary Figure 2-1)**. Otherwise, the distributions of reaction times appeared broadly comparable.

For hand responses, monkeys could use either or both paws on the touch screen, while humans used the fingers of the preferred hand for button presses. Nevertheless, the monkeys were also somewhat faster than the humans on the RDK task, when responding by hand (mean RT = 0.631s, STD = ± 0.215s vs. mean RT = 0.724s, STD = ± 0.355s; independent t-test, *p* < 0.0001). A closer look at the reaction times across different stimulus levels shows the typical pattern of longer reaction times for stimuli that are more difficult to judge (**Supplementary Figure 2-1**). This difference was more pronounced for the RDK than for the flatter cylinder reaction times, especially for the human data. Notable is also m133 who showed a bias in the RDK task with different reaction times for right and left choices, which most likely indicates a hand bias.

### Visual stimulus but not response mode produces systematic differences in drift constant and boundary separation in drift diffusion modelling

We constructed hierarchical DDM models for the pooled data from the 20 human participants across all four task combinations and for each macaque separately across the two available task combinations, yielding a total of 8 fitted models (four fits from the 20 humans and two fits from each monkey). To compare performance across the two different tasks with different visual stimuli, human psychometric data sets were normalized by dividing each stimulus value by the average threshold of the group of human participants for this task. For the monkey data, we normalized by their individual thresholds across sessions. We utilized the open-source Bayesian hierarchical DDM method (Wiecki et al., 2013), which assumes the individual data sets come from a population distribution and estimates the model parameters using the Monte Carlo Markov Chain (MCMC) method. We treated each human participant as a member of a population and, similarly, all experimental sessions for each monkey on a given task as members of a population. This latter approach was used due to the small number of monkeys (Wiecki et al., 2013). The models were constructed with 5 parameters: boundary separation *a*, drift constant (a form of sensitivity) *k*, drift intercept *v*_0_, non-decision time *t*, and starting point bias *z* (see **Figure 1C**). We ran the Monte Carlo Markov Chain (MCMC) of the DDM five separate times for each model to assess convergence as well as for further analysis. The Gelman-Rubin statistic (Gelman & Rubin, 1992) was calculated to lie between 0.99 and 1.01 in all cases except one (1.013), showing excellent convergence.

A pair-wise comparison between the 3D cylinder task and the RDK task of the distribution of fitted DDM parameters reveals systematic differences for the drift constant *k* and for the boundary separation *a* (**Figure 3**, panels surrounded by dashed black boxes) (Kolmogorov-Smirnov test p < 0.0001 (test statistic 0.825) for drift constant *k*, p < 0.001 (test statistic 0.45) for boundary separation *a*). This is the case for humans and monkeys, regardless of the mode of response. For the goodness-of-fit see **Supplementary Figure 3-1**. The value of *a* was lower in the cylinder task than for the RDK task in both species, indicating a shorter distance to evidence threshold. The human participants had a larger drift constant *k* for the RDK tasks, suggesting that the RDK sensory signal was integrated into evidence at a steeper rate than the disparity information. While the monkeys also showed a strong difference in drift constant *k* between the cylinder and the RDK task, the monkeys showed a larger drift constant *k* for the cylinder task.

**Figure 3.**
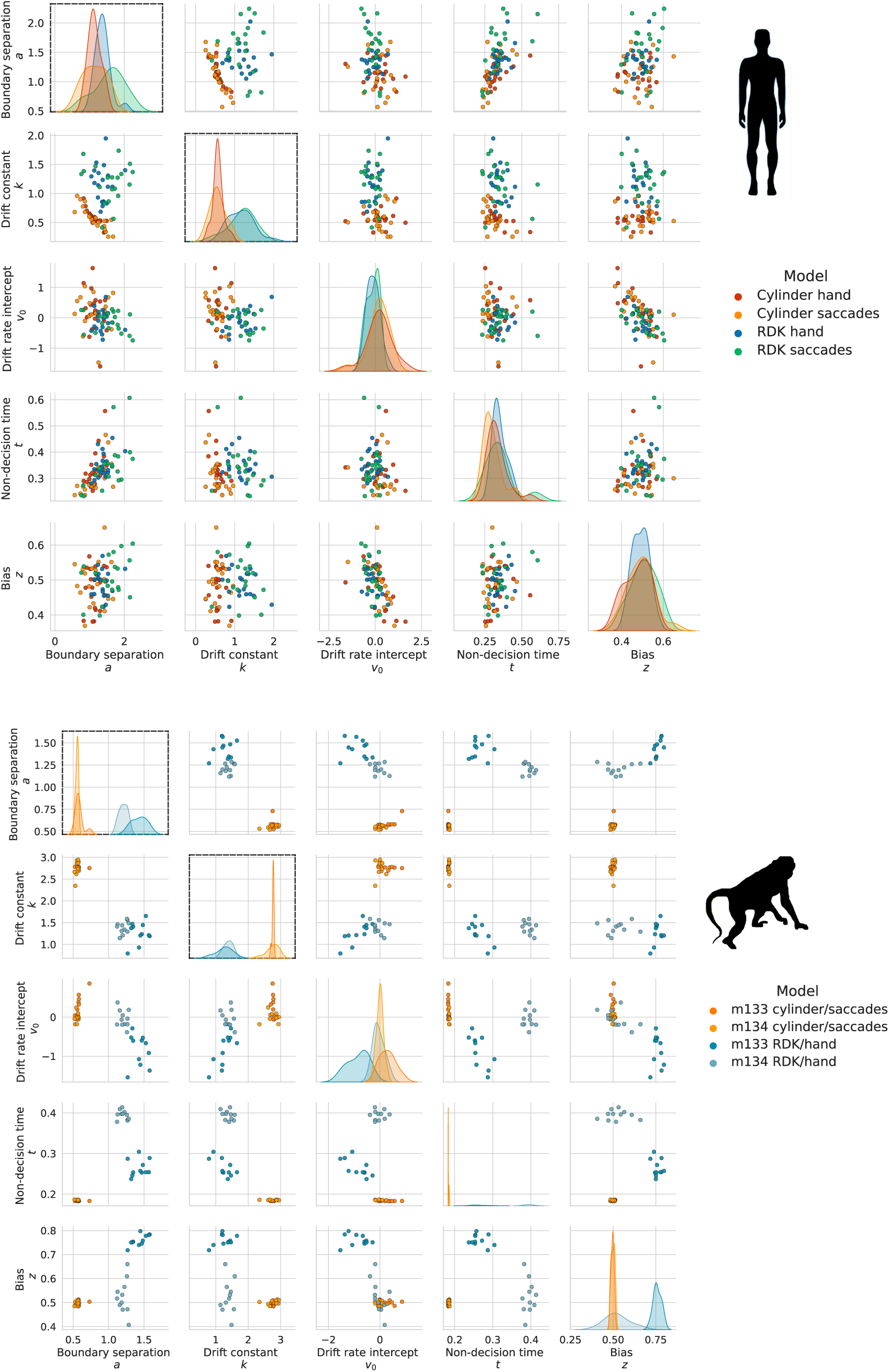
Pair-wise comparison of estimated parameters for the DDM. DDM Parameters are shown separately for humans (n = 20, upper panel) and monkeys (n = 2, bottom panel) as scatterplots with kernel density estimates on the diagonal. There are statistically significant differences (panels with dashed black outlines) between the parameter distributions for boundary separation *a* and drift constant *k*, for both monkeys and humans, between the different stimulus types (3D SfM cylinder vs RDK motion—see main text for statistics). The boundary separation *a* is lower for the cylinder rotation task determined by binocular depth than for the direction discrimination task with the RDK in both species, regardless of the type of motor response. The drift constant *k* also differs systematically for the two stimuli. It is higher for humans judging RDK motion direction. There is also a clear difference for the monkeys but here the pattern is reversed with the cylinder task showing a higher drift constant *k*. There is no systematic difference with response mode. The monkey data also reflect some of the individual differences between the animals, for instance m133 showed different reaction times when responding left or right on the touch screen (see **Supplementary Figure 3-1**). This is reflected in the bias *z*.

Because the monkey data were obtained from only two subjects, individual differences are more apparent in the data set. In contrast, the human data were obtained from 20 individuals for which the parameter values are more spread out and appear normally distributed. Nevertheless, the data clearly indicate systematic differences in two key DDM parameters, *k* and *a*, across both species. These differences depend on the type of visual stimulus (binocular depth defined rotation or direction of motion) about which decisions are made, regardless of the mode of response (hand or eye).

### Dimensionality reduction confirms a systematic difference by visual task

To tease apart the factors contributing to the differences in DDM parameters for humans and monkeys, we implemented a supervised dimensionality reduction method, linear discriminant analysis (LDA), on the DDM model parameters shown in **Figure 3**. Using the estimated parameter distribution derived from the 5 runs of the MCMC model on the human data, we reduced the space to three linear discriminant dimensions (LD) which optimally distinguished the four groups (combinations of stimulus type and response mode). We visualized this by projecting the human estimated parameters onto the first two LDs. Subsequently, we projected the estimated parameters for the monkey, also combined from 5 MCMC runs, onto this same space derived from the analysis of the human parameters (**Figure 4**).

**Figure 4.**
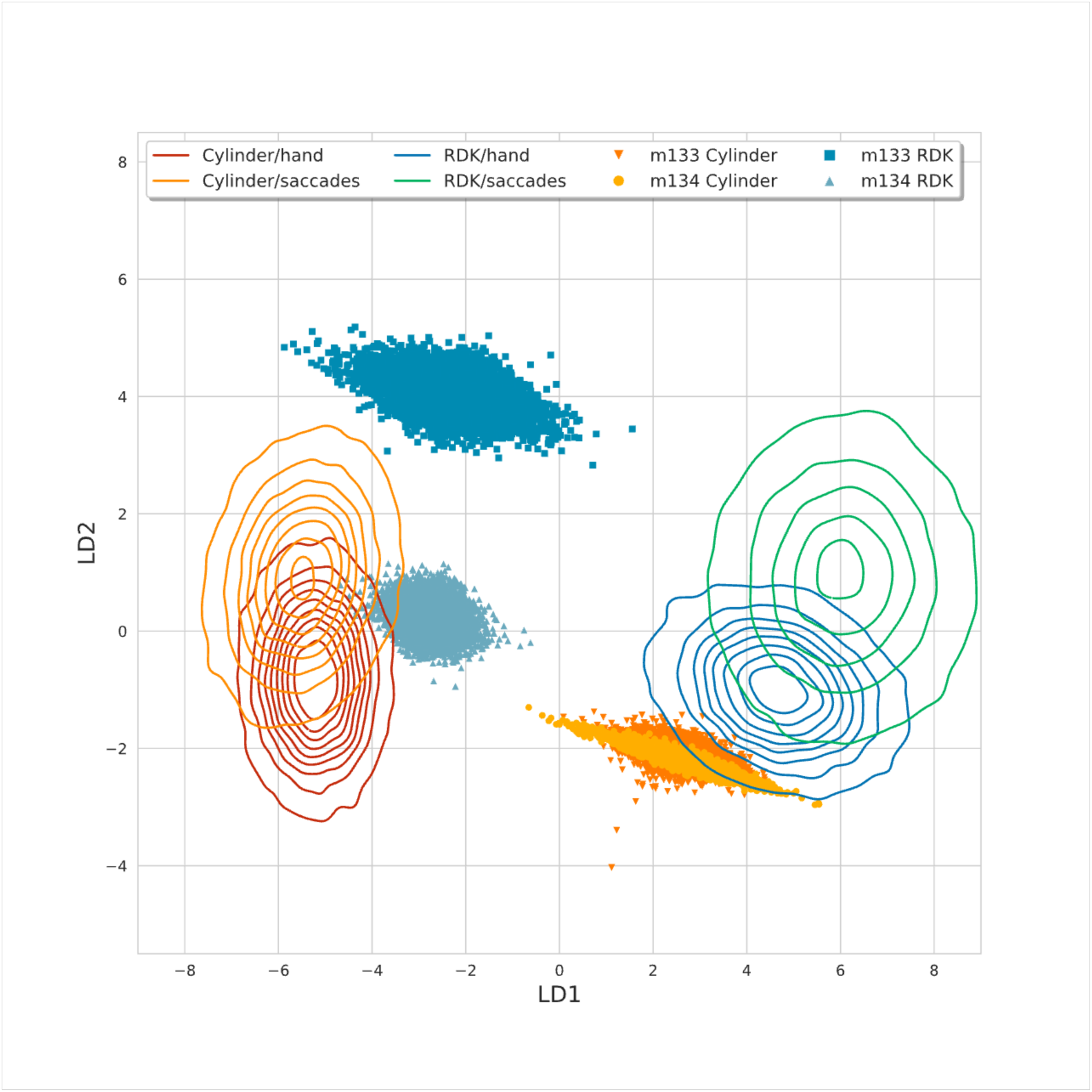
Linear discriminant analysis (LDA) confirms differences with visual stimulus type. The first two linear discriminant dimensions of the LDA for the estimated DDM parameters were plotted along the abscissa and ordinate, respectively. The estimated parameters for the human data are depicted as contour plots (representing iso-proportions of probability density), red and orange for the 3D SfM cylinder task and blue and green for the RDK direction discrimination task. LD1 clearly discriminates the clusters by stimulus type, with a smaller difference but considerable overlap on LD2 by response mode. The estimated parameters for the monkey data were projected in the form of scatter points onto this space by transforming it with the vectors derived using the human data. The two monkeys also show along LD1 a clear segregation of the DDM parameters by stimulus type about which perceptual decisions are made, although with a sign reversal to the humans. They show some differentiation by response mode along LD2, but with sizeable individual differences.

Immediately apparent are the separate clusters along LD1 for the two visual stimuli (RDK motion direction or cylinder). Along LD2, mode of response was a poor separator for the human data, but somewhat better for the monkeys. Each LD was a weighted combination of the 5 original DDM parameters. For LD1, which shows the highest discriminability, *k* contributed 69%, *a* 17%, *v*_0_ 6% and whereas *t* and *z* showed negligible contribution (less than 5 %). For LD2 *k* contributed 26%, *a* 34%, *t* 21%, *z* 19% and *v*_0_ showed virtually no contribution. The DDM parameter space is therefore distinct for perceptual decisions about the RDK stimulus and the rotating cylinder. Overall, the DDM suggests distinct decision processing for the two different visual stimuli, but not for the saccade or hand response. The estimated parameters for the monkey data are largely comparable in this respect with those for the human data, except for m133 on the RDK task, for which the monkey showed a bias in the RT for different directions.

### Hidden Markov Model reveals different strategies for the two visual tasks

Looking at choice only, we probed whether the observed differences in decision-making between the two visual tasks were related to distinct behavioural strategies when judging the two visual stimuli. We employed a generalized linear model (GLM) with a logistic function as the link function, translating the sensory input into the probability of choosing “rightwards direction” (RDK) or “counter-clockwise rotation” (cylinder), and 3 covariates, namely normalised *stimulus* strength, previous *choice*, and *bias* (Ashwood et al., 2022). The human data were grouped by the type of stimulus to be judged. We then modelled a number of distinct “behavioural states” as different states in a Hidden Markov model (HMM) by estimating weights for each of the covariates using an expectation-maximization algorithm with 5-fold cross-validation for humans and *K*-fold for monkeys depending on the number of sessions *K* (Ashwood et al., 2022).

We compared the rise in log-likelihood (LL) per trial averaged across the five hold-out sets. Each behavioural state is described by a distinct set of weights attributed to the three covariates, which describe the behavioural strategy of the animal on these trials (**Figure 5**). By identifying the steepest rise in LL, we found that three states were optimal to describe the human data for each stimulus type (**Supplementary Figure 5-1**). Based on the pattern of weights for the two most distinguishing covariates, *stimulus* (indicating the task engagement by linking stimulus strength to choice) and *bias*, these states were *post-hoc* termed “Strongly engaged”, “Moderately engaged with bias” and “Disengaged” for the 3D SfM cylinder and “Engaged”, “Engaged with left bias” and “Disengaged” for the RDK task (**Figure 5A**).

**Figure 5.**
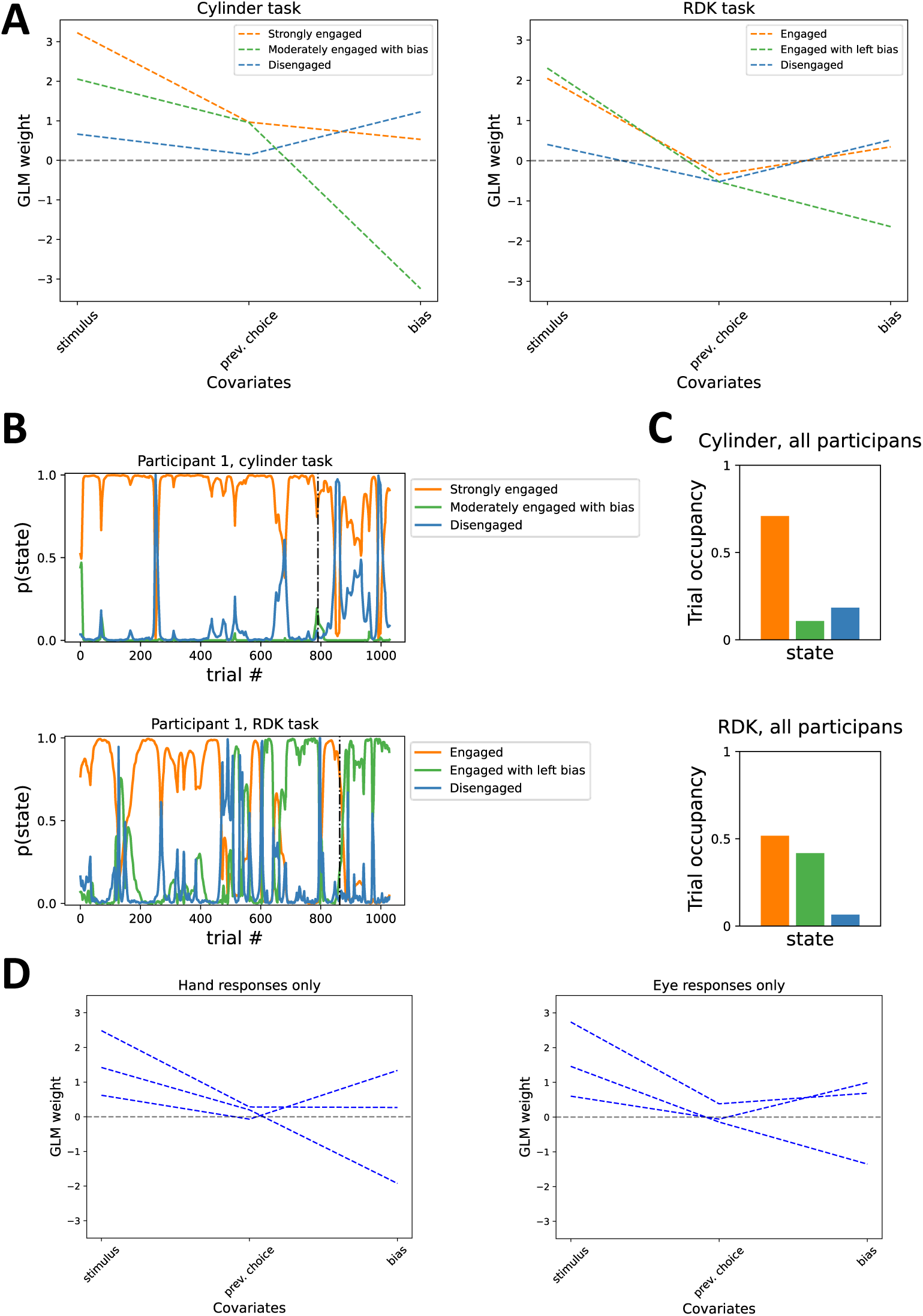
The GLM-HMM analysis reveals distinct behavioral states for perceptual decisions. **A**. The global fits for the human participants (n = 20) show three distinct states for each stimulus type. The three states between which the participants switch vary for judgements about RDK direction of motion and 3D SfM cylinders, especially in the engagement level (influence of the stimulus) and the direction of the effect of previous choices on the current choice. **B**. The posterior probability of being in each state for an example participant performing the cylinder and the RDK task fluctuates more between the three states for the RDK stimulus than for the SfM cylinder (see **Supplementary Figures 5-3, 5-4** for all individual participants). The dash-dotted line separates trials with the hand responses on the left side and the saccadic response on the right side (see Methods). **C**. The fractional occupancies, or the proportion of trials in which the posterior probability was highest for a certain state, show that for the cylinder stimulus, human participants spent most of the time in a strongly engaged state without much evidence for a behavioural or perceptual bias. In contrast, for judgements of the RDK stimulus, humans switched between two engaged states, one with a strong bias influencing perceptual decision-making. **D**. When the human data was grouped for the mode of response, the three states appeared qualitatively very similar between the hand- and eye-responses.

The 3D cylinder showed the overall strongest task engagement in the “Strongly engaged” state. Participants spent the majority of their time in this state (**Figure 5B top, 5C top**). In contrast, participants judging the RDK spent roughly equal time in two “engaged” states with an influence of the *stimulus* on choices, which was similar to that found for the “moderately engaged state with bias” for the cylinder (**Figure 5C bottom**). One of these two RDK engaged states showed a bias, but not the other. When judging the 3D cylinder, human participants spent considerably more time in an engaged state without a behavioural bias than for discriminating motion direction of the RDK. Finally, for both stimuli, we identified a “disengaged state”, during which *stimulus* only weakly controlled the responses.

The covariate *previous choice* seemed to have a smaller influence on the individual states, but there was a pattern of 3D cylinder states tending to stay with a previous choice (*previous choice* +1), while RDK states tended to favour switch away from the previous choice (*previous choice* -0.5). When we re-grouped the human data according to the mode of response, again three states were sufficient. However, the states appeared almost indistinguishable between responses with eye or hand movements (**Figure 5D**).

For the monkey data, grouped individually by animal and task, the LL suggested that two states were slightly better for both tasks for each animal (**Supplementary Figure 5-2**). This conclusion could also supported for a small number of the human participants, based on further analysing the individual trial occupancies (**Supplementary Figures 5-3** and **5-4**). For the monkeys, the identified states corresponded broadly to a subset of those found in humans for the same tasks, particularly the fit for m134 was very similar to the global fit for the human data (see **Supplementary Figure 5-2 for states** and **Supplementary Figures 5-5** and **5-6** for the individual trial resolved data).

Overall, the analysis of behavioural strategy for the human participants complemented the findings of the DDM, in that the two visual stimuli did exhibit different state distributions, whereas changes of state did not associate with the two modes of response. It also emerged that different periods of time were spent in distinct behavioural states when judgements were made about the two types of visual stimuli and there was some similarity in this regard between the humans and the monkeys

In order to test whether both modelling approaches pick up consistently on the same differences between the two visual tasks, we identified for the humans the GLM-HMM states with the largest number of trials (“3D cylinder: strongly engaged”, n=17 participants, “RDK: moderately engaged” n=12, and “RDK: moderately engaged with left bias” n=7) and performed a second DDM run, including data from participants with a minimum number of contributing trials. When we compared the DDM parameters for these three states, they also showed the distinct separation in drift constant *k* and boundary separation *a* between the two stimulus types (**Supplementary Figure 5-7).** This suggests that the two modelling approaches pick up on corresponding differences in processing. The behavioural state for the RDK influenced by a bias shows a strong drift rate intercept.

In summary, both models distinguish consistently behavioural differences in judging the two visual stimuli but less so or not all for the modes of response. The DDM suggests differences in sensitivity (*k*) to the sensory information and time available for the decision process (*a*). The GLM-HMM also points to distinct behavioural strategies or brain states for decisions about each of the two types of visual stimuli. These differences might be related to the strong, stable percept that subsumes all stimulus elements (dots) of the cylinder stimulus, even in the bistable version. In contrast, decisions about the more overtly noisy RDK stimulus need to discount the noise dots and the perceptual process might be consequently less stable. Ultimately, the differences in perception point to differences in the brain processes responsible for perceptual decision-making in the two tasks.

## Discussion

We systematically analysed perceptual decisions in a multi-factorial psychophysical design for combinations of two distinct visual cues to be judged with two modes of perceptual report by humans and macaque monkeys. These cues of motion and binocular disparity were presented in the form of stimuli (RDK and 3-D cylinders) that have been extensively studied electrophysiologically and causally linked through microstimulation to macaque area V5/MT. The two analyses (DDM and GLM-HMM) offered complementary and consistent findings.

Firstly, we tested whether the computational models could underpin distinct processing for the two visual tasks and the different motor responses. Both the DDM and GLM-HMM identified clear separation in the parameter space of the fitted models for judgements of motion direction in RDK versus perceived rotation of 3D SfM cylinders. Conversely, for the type of motor response, there was only a weak difference in the DDM and none in the GLM-HMM. Secondly, we set out to compare humans and monkeys in the same modelling frameworks. Like humans, the monkeys showed a clear distinction between the two visual decision tasks. However, some of the details of the fitted models were different on this small sample. We propose that perceptual decisions might be affected by different integration windows for incoming visual information about motion and about 3D depth.

### HDDM and GLM-HMM point to different neuronal mechanisms and behavioural strategies for the two visual tasks

The DDM parameter space showed a clear separation of the models for the different visual stimulus cues. For the human as well as the monkey data, the separation of the bounds (*a*) was lower for the cylinder than for the RDK. In the human data, this was also accompanied by lower drift constant (*k*) for the cylinder stimulus. This implies that a shorter integration window was needed to commit to clockwise or counter-clockwise rotation in comparison to the time needed for decisions about left- or rightwards motion in the RDK. The outcome is most evident for the more ambiguous versions of the stimuli (see **Supplementary Figure 2-1**). At the same time, the lower drift constant for the cylinder task compared with that for the RDK task in the human data suggests that binocular depth matching is a fundamentally more difficult task in terms of translating the signal into evidence.

In the decision task, the bottom-up visual input leads to the selective activation of the relevantly tuned neuronal populations in visual area V5/MT for both stimulus types (Britten et al., 1992; Dodd et al., 2001; Krug, 2020). In the case of the 3D cylinder, we propose that this would activate circuits whose positive feedback activity in the sensory cortex stabilizes the network of neurons favouring one direction of rotation over the other. Such reverberations could theoretically stabilize the network faster than for the corresponding case with the RDK, leading to lower effective separation of the bounds.

The reaction times on more difficult trials are almost as fast for the cylinder as the easiest trials with a strong signal – in contrast to the RDK RTs. This may also be attributable to greater positive feedback within the local network dynamics during perceptual decisions about the cylinder. The two monkeys, however, showed the opposite tendency: The drift constant for monkeys for SfM cylinders is higher than for the RDK stimulus. This outcome might be attributable to perceptual learning, given that the macaques underwent far more extensive training on the cylinder task than the humans prior to data collection and learned the cylinder task after the initial training on the RDK task. Further data are needed, preferably conducted with electrophysiological recordings during the training period.

The GLM-HMM matched choice data to hypothesized “brain states”, which are manifest as different weights for the parameters that contribute to decision-making processes. These parameters include the stimulus to be judged, the choice made on the preceding trial, and internal biases. When the human data were grouped according to the motor response, the states identified for hand and eye responses came out very similar in terms of their GLM weights. Clear differences emerged when the same data were grouped based on the visual stimulus to be judged. Most striking, when judging the SfM cylinder, human participants tended to stay in one “strongly engaged” state far more than in the other two states (“moderately engaged with a bias” and “disengaged”).

By comparison, when deciding about the motion direction of an RDK, human participants tended to switch between two distinct “engaged” states (one of them with a strong bias). We see evidence of this not only from the fractional occupancy of the three states but also in the trial-resolved data (see **Figure 5**). Moreover, weighting of the previous choice differed in sign between all the RDK states (negative) and the cylinder states (positive), which means the participants tended to switch from their previous choice when viewing RDKs but tended to stay with their previous choice when viewing cylinders. These observations might be potentially be ascribed to features of the neuronal network underpinning the cylinder task which might make it harder to “snap out” of a particular state, for example to a stronger local feedback loop within the neuronal circuitry supporting a stable 3D cylinder percept. Conversely, the behaviour of the RDK-subserving network might be better explained by sensory adaptation carrying over from one trial to the next.

### Potential neural substrate and mechanisms

For both perceptual decision tasks, extrastriate visual area V5/MT is firmly implicated as a potential neural substrate, based on a robust experimental framework linking single neurons to perception (Barlow, 1972; Parker & Newsome, 1998). Electrical microstimulation of focal sites in V5/MT cortex alters direction of motion and 3D SfM cylinder percepts in a predictable and constructive way (Salzman et. al., 1990; Krug et al., 2013). Single V5/MT neurons of macaque monkeys show steep neurometric functions for the RDK task, matching those of the psychometric functions of the animals (Britten et al., 1992; Newsome et al., 1989). An above-chance relationship was also established between the trial-by-trial fluctuations in the activity of the V5/MT neurons and the perceptual decisions made by the animal about RDK stimulus, independent of the strength of visual stimulation; this relationship is termed choice probability (CP) (Britten et al., 1996). CP is defined as the probability with which an independent observer could predict the choices of the animal from the neuronal activity alone, given knowledge of the tuning curves.

CPs above chance level for different tasks have been found across extrastriate visual areas (Nienborg et al., 2012), with two broad interpretations with regards to their origin: bottom-up and top-down input. The former can be formalised as a feedforward neural network model wherein the final choice is determined by comparing two pools of activity of neurons with opposite tuning and, the activity within the pools is correlated (Shadlen et al., 1996), see also (Haefner et al., 2016). However, experimental data apparently at odds with this model have been obtained, giving rise to the latter explanation, that top-down control from downstream areas contributes to shared variability of activity, which is revealed as raised levels of interneuronal correlation (Nienborg & Cumming, 2009). Experimentally derived CPs and interneuronal correlations from visual area V5/MT of macaque monkeys making perceptual decisions about the 3D SfM cylinder were both higher (CP = 0.67, interneuronal correlation r = 0.37) (Dodd et al., 2001; Wasmuht et al., 2019)(Krug et al., 2016) than those reported for the RDK task (CP = 0.56, r = 0.17) (Bair et al., 2001; Britten et al., 1996; Zohary, Shadlen, et al., 1994). These data suggest differences in neuronal processing within the local circuitry of V5/MT may underlie the psychophysical differences found here for the two stimuli.

Previous studies have investigated the network dynamics of V5/MT, both experimentally as well as theoretically. Wasmuht and colleagues (2019) concluded that the interneuronal correlations underlying the CP were dependent on interactions at longer timescales (100 – 400 ms) for ambiguous cylinders than RDKs, suggesting a stronger feedback component to the emergence of CP. A recent computational model accounts for the sustained CP observed experimentally whereby a bottom-up component of the interneuronal correlations explains the initial period of the decision time and a top-down component contributes to a second, later time period (Wimmer et al., 2015). However, if this top-down signal contributes to decision-formation, this would break a fundamental assumption of the DDM model, since this model accumulates only external evidence to form the decision. Nevertheless, one prediction of such a feedback or top-down mechanism is a shorter effective information integration window during which the bottom-up component can most effectively steer the decision process. This should lead to a flatter distribution of RT as a function of difficulty in the 3D cylinder task compared to the RDK task (Wimmer et al., 2015), as we observed in **Supplementary Figure 2-1.**

In the cylinder task, electrophysiological measures show that interneuronal correlation is weaker when the bottom-up task-relevant input, the magnitude of binocular disparity in the stimulus, is greatest. The strongest interneuronal correlations arise for the ambiguous cylinder, when the top-down decision signal may be at its strongest. Wasmuht and colleagues (2019) also showed that CP from V5/MT of macaque monkeys performing the cylinder task was positively correlated with interneuronal correlations and specifically associated with their long timescale component. This finding is consistent with the interpretation that feedback connections establish an increase in interneuronal correlations and a rise in CP in V5/MT, resulting in a stabilization of the percept. For the RDK stimulus with the weaker perceptual coherence, weaker CPs and interneuronal correlation suggested that the feedback or top-down modulation was not nearly as powerful. We suggest that the SfM cylinder task involves network dynamics with more rapidly rising and stronger top-down feedback, which consequently generates higher interneuronal correlations between neurons with similar preference as well as higher CPs.

About the mechanisms that result in a shorter integration window within the same neuronal circuitry, we can, at present, only speculate. It is clearly essential to perform direct electrophysiological measurements of multiple single neurons during perceptual decision-making with both the cylinder and the RDK tasks. However, we tentatively offer the following explanation. It has been suggested in (Dodd et al., 2001) that the percept of the ambiguous cylinder rotating in one direction is very different from the same cylinder rotating in the other direction, but the percept of the ambiguous RDK with one selected direction of global motion is much more similar to the percept of the same RDK with a choice in the other direction. In this interpretation, the two perceptual decisions are much farther apart in neuronal feature space for the case of the cylinder compared with the RDK. This greater separation in feature space may be linked to the presence of two separate but synergistic processes that must be recruited in the cylinder task, namely depth-from-motion (kinetic depth) and depth-from-disparity (dynamic stereopsis) (Nawrot & Blake, 1993).

## Conclusion

Computational models in systems neuroscience provide a powerful and robust framework for predicting neural mechanisms underlying cognitive processes. We systematically investigated two perceptual decision tasks with two distinct motor reports using the established DDM and the more recent GLM-HMM and interpreted our results in the context of a large body of existing electrophysiological literature. Both models clearly showed distinct decision processing for the two visual cues, direction of motion and 3D depth, but not for the two modes of response, hand or eye movement. The results were broadly similar between humans and monkeys on these tasks. We propose that these results reflect distinct processing for decision-making about the two stimulus types, with a shorter integration window for bottom-up input for 3D signals in the SFM cylinder than for motion direction in the RDK stimulus. To understand these mechanisms underlying perceptual decisions at a biologically plausible level of description across primates, the next steps should be to combine multisite neurophysiological recordings with models of neural dynamics at the level of microcircuits (Hanks & Summerfield, 2017). The circuits of extrastriate visual area V5/MT with its columnar architecture for direction of motion and 3D depth and specific long-range interconnectivity are a likely substrate (Ahmed et al., 2012; DeAngelis & Newsome, 1999).

## Materials and methods

### Human participants

We collected data from 23 adults of which 20 completed the study (13 females, 7 males; aged between 18 and 40, mean age of 29.6 years). Except one female, all were right-handed. All participants passed a test for normal or corrected-to-normal visual acuity (Snellen chart with 6/9 or better). Additionally, all participants passed a test of static stereoscopic vision (TNO, Lameris Ootech) using random-dots stereograms and red-green glasses with a minimum threshold of 240 arc seconds. Ethical approval was obtained from the Coordination Center for clinical studies (KKS) at the Medical Faculty of the Otto-von-Guericke University Magdeburg (OVGU). The study was conducted in accordance with the data protection principles (Datenschutz-Grundverordnung: DS-GVO) and the Declaration of Helsinki (2013). All participants gave informed consent.

### Animals

We collected data from two male adult rhesus macaques, m133 and m134. Each animal was first trained on the RDK task with a touchscreen and then subsequently on the cylinder task while head-fixed and responding through saccades. All procedures were conducted under licenses from the United Kingdom Home Office in accordance with the Animals (Scientific Procedures) Act 1986 and the European Union guidelines (EU Directive 2010/63/EU).

### Human experimental set-up and visual stimuli

Data were collected at a psychophysical laboratory at the Institute for Biology of the OVGU. The laboratory was darkened with blinds and shutters blocking all external light and additionally with black-painted dividers around the visual set-up. Visual stimuli were presented using a Wheatstone stereoscope on two EIZO FlexScan L550 monitors (refresh rate 60 Hz, 338 mm width, 270 mm height, resolution of 1280 by 1024 pixels) at viewing distance of 56 cm. Anti-aliasing was implemented to decrease the minimal effective disparity that could be displayed. Participants indicated their decisions using either two buttons on a response box or by making a saccade to a visual target. Eye movements were tracked throughout experimental trials using EyeLink 1000 Plus (SR Research EyeLink, Canada) at 1000 Hz sampling rate.

Visual stimuli were coded in MATLAB using Psychtoolbox (Brainard, 1997) and displayed on mid-grey background. The circular RDK patch had a diameter 6° and was made up of 180 black and white dots. Dot diameter was 0.1° and moved at the speed of 3°/s. Dots moved in random directions with a given percentage moving either left or right on a given trial, referred to as coherence. All human participants viewed the same coherence levels in a randomised order: -50%, -25%, -12.5%, -6.25%, - 3.125%, 0 %, +3.125%, +6.25%, +12.5%, +25%, +50% (the sign indicates opposite directions). The 3D SfM cylinder is a projection of a 3D cylinder on a 2D screen (Dodd et al., 2001). Like the RDK, it was also made up of 180 black and white dots in on a mid-grey background. Dots were distributed on two superimposed, transparent planes. Dots on the two planes moved in opposite directions (right or left) with as sinusoidal velocity profile. Cylinder size was 6° by 6° and dot diameter 0.1°. Binocular disparity applied to the two sets of dots was used to separate the two transparent planes and disambiguate the cylinder rotation direction. Cylinder disparities were - 0.08°, -0.04°, -0.02°, -0.013°, -0.007°, 0°, +0.007°, +0.013°, +0.02°, +0.04°, +0.08° between the centre of the back and front surface, randomly interleaved. To prevent dot tracking, 1 % of the dots was removed and replaced every frame (16.6 ms) for both stimuli.

For each participant, experimental sessions were spread over four separate days, most participants completed all four sessions within two weeks. Each session consisted of eight experimental blocks of 110 trials each with a 2-minute mandatory break between each block; longer breaks were taken as requested by participants. At the beginning of each session, participants were given training blocks to ensure stable performance, with their threshold being lower than at least the largest disparity presented on those training blocks. A typical experimental session lasted 1.5 – 2 hours. Participants conducted the study in 1 of 4 possible combinations with different stimuli and responses, listed in **Table 1**. We employed a balanced design and the assignment to groups was random.

**Table 1.** The four experimental orders the human participants were randomly assigned to, forming balanced groups of 5. C = Cylinder; R = RDK; S = Saccades; B = Buttons.

| Experimental order | Sequence of sessions |
| --- | --- |
| 1 | CS, RS, CB, RB |
| 2 | CB, RB, CS, RS |
| 3 | RS, CS, RB, CB |
| 4 | RB, CB, RS, CS |

At the beginning of each session, participants were briefed on the set-up and the task. To start each trial, participants fixated on a central dot (radius 0.13°) flashing black and white at 5 Hz, which then turned white. They were required to maintain fixation within a 1.7-degree radius around the fixation point. After acquisition of the fixation point, the stimulus appeared in the centre of the screen. Participants were instructed to report the direction of motion or the direction of rotation as fast and as accurate as possible. In sessions when an eye movement response was required, two small saccade targets (diameter = 0.5°) were displayed 6° to the left and the right on each side of the fixation dot. When a hand movement response was required, participants pressed with one of two buttons (**Figure 1A**). Participants had to indicate their choice within 3s, otherwise the trial was aborted. Auditory feedback was given with a high-pitch tone for correct choices and a low-pitch tone for incorrect choices. Randomly one of the two tones was played for responses to ambiguous stimuli. The inter-trial interval was 1.5 seconds.

### Monkey experimental set-up and visual stimuli

Both monkeys underwent first training on the RDK task. For this study, they performed 11 consecutive sessions, each on a different day, within the span of 3 weeks. Monkeys were positioned in a monkey chair in front of a touchscreen (Elo Touch Solutions, LCD Touch Monitor, 60 Hz, Leuven, Belgium) with a viewing distance of 20 cm. Visual stimuli were coded in MATLAB using Psychtoolbox (Brainard, 1997) and were displayed on mid-grey background. Each trial started when the monkey touched a flashing dot on the bottom half of the screen. Then the RDK appeared above the dot together with two white targets in the form of a circle and a cross (diameter = 6.75°) to the left and the right. The RDK patch (diameter = 17.38°) was composed of 150 black and white dots (dot diameter = 0.5°). Dots moved in random directions with a certain percentage coherently moving left or right at a speed of 17.38°/s. m133 judged motion coherence levels of -50%, -40%, -30%, -25%, -20%, -12.5%, -10%, -6.25%, -3.125%, 0%, +3.125%, +6.25%, +10%, +12.5%, +20%, +25%, +30%, +40%, +50% (the sign indicates opposite directions). m134 judged coherence levels of 50%, -25%, -12.5%, -6.25%, -3.125%, 0%, +3.125%, +6.25%, +12.5%, +25%, +50%. Monkeys were rewarded with juice when they touched the correct target corresponding to the global motion direction. They were rewarded at random for ambiguous stimuli.

Subsequently, the same two monkeys underwent training for the 3D SfM cylinder paradigm. Monkeys were head-fixed during this task with an implanted titanium headpost. Visual stimuli were presented using a Wheatstone stereoscope on two EIZO FlexScan F78 CRT monitors (refresh rate 85 Hz, 392 mm width, 294 mm height, resolution of 1600 by 1200 pixels) at a viewing distance of 84 cm. Eye movements were tracked using the IVIEW X Hi-Speed Primate Eye Tracker (SensoMotoric Instruments SMI, Germany). Each trial was started with the monkey fixating on a dot (radius 0.13°) located -2° down from the centre of the screen. Fixation within a radius of 1.7° rendered the black and white at 5 Hz flashing fixation dot white and the cylinder appeared 5° vertically above it. Simultaneously two white choice targets appeared on each side +4.5° from the cylinder stimulus in the form of a circle and a cross (diameter = 1.5°). The cylinder was made up of 180 black and white dots placed on two transparent planes. The cylinder size was 6° by 6° and the dot diameter 0.2°. The cylinder disparities used to disambiguate the direction of rotation were: -0.03°, -0.02°, -0.015°, -0.01°, -0.005°, 0.00°, +0.005°, +0.01°, +0.015°, +0.02°, +0.03°. To prevent dot tracking, the mean dot lifetime was 45 frames (750 ms). Monkeys reported the rotation percept with a saccade to one of the two choice targets within a 2 second window. Correct trials were rewarded with juice. For ambiguous trials, only about half of the trials were rewarded. This was unintentionally only the rightward choices. Because the ambiguous trials are embedded within a large set of sub-threshold trials and are not distinguishable from these or even near-threshold trials (Nawrot & Blake, 1993), this did not lead to a bias in behaviour. Overall, both monkeys made only about 55% rightward choices on ambiguous trials.

### Behavioural data and analysis

We implemented outlier removal for the reaction time data, as is common practice when fitting DDM and other models (Berger & Kiefer, 2021; Ratcliff & McKoon, 2008). The cut-off points were based on pre-existing literature as well as examination of the behavioural performance within certain time intervals, such as 150 – 190 ms (**Supplementary figures 2-2 and 2-3**; **Table 2**). Furthermore, 5% outlier removal was implemented within these boundaries in the HDDM fitting procedure to exclude the trials least likely to have been generated by a sequential sampling process for each model (Ratcliff & McKoon, 2008).

**Table 2.**
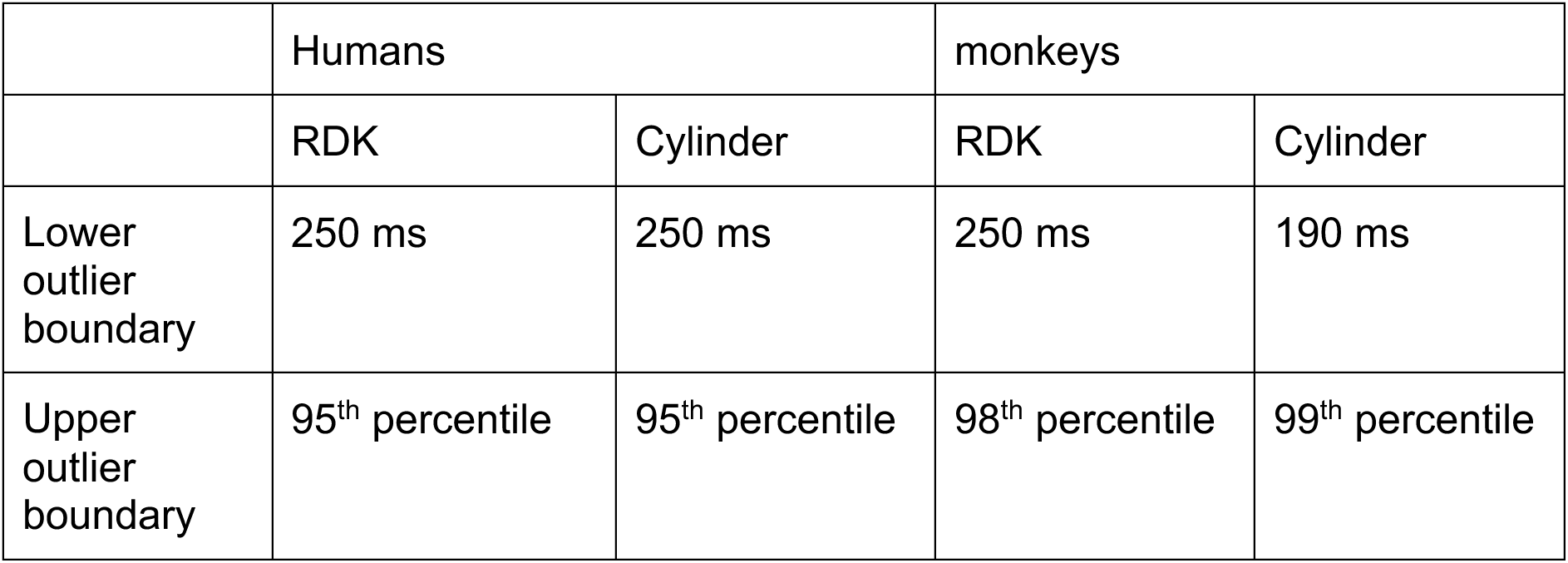
The outlier removal used in data pre-processing. The 98^th^ and 99^th^ percentile as the upper outlier boundaries were chosen to best capture the bulk of the distribution in each case since the distributions differed in their skewness. Because the trials of the cylinder task for each monkey were more numerous as well as more compactly distributed (with a shorter right tail), we opted to include one more percentile than for the RDK data.

|  | Humans |  | monkeys |  |
| --- | --- | --- | --- | --- |
|  | RDK | Cylinder | RDK | Cylinder |
| Lower outlier boundary | 250 ms | 250 ms | 250 ms | 190 ms |
| Upper outlier boundary | 95<sup>th</sup> percentile | 95<sup>th</sup> percentile | 98<sup>th</sup> percentile | 99<sup>th</sup> percentile |

Psychometric curves (**Figure 2**) were fitted according to **Equation 1**, describing a cumulative Gaussian distribution, where *x* is the level of signal, for example binocular disparity in degrees, *μ* is the mean of the distribution, equivalent to the bias, and *σ* is the standard deviation of the distribution, equivalent to the threshold. We used a nonlinear least-squares solver implemented in MATLAB (*lsqcurvefit*), which minimises the sum of squared errors, to estimate the two parameters.

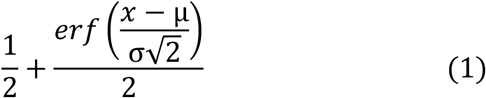

### DDM fitting

We used the hierarchical DDM (HDDM), a python toolbox for hierarchical Bayesian parameter estimation, to fit the model to our data (Wiecki et al., 2013). The HDDM allows simultaneous estimation of subject and group parameters, where individual subjects are assumed to be drawn from a group distribution. There are a number of different implementations of the DDM fitting frameworks, broadly comparable in performance (Ratcliff & Childers, 2015; Shinn et al., 2020). HDDM was chosen as it implements a Bayesian framework and estimates distribution of parameters instead of point estimates allowing better exploration of parameter space and statistical analysis of parameters. The HDDM was also a good design for our multi-condition experiments across individual subjects due to its suitability for estimating parameters with fewer data points.

In these models, a time-dependent decision variable (*DV*) is driven by the sensory samples and represents the current amount of available evidence. The value of *DV* is described by a Wiener diffusion process, in which *DV* evolves via the stochastic differential equation 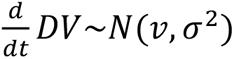 where *v* is the drift rate. The marginal distribution over *DV* at any given time *t* is described by *N*(*vt*, *σ*^2^*t*). The choices and corresponding RTs are treated as random variables described by the first passage times for this Wiener process (Navarro & Fuss, 2009). We can expand the drift rate parameter to its intercept *v*_0_and the scaling factor *k* which translates the signed normalised evidence signal strength according to **Equation 2**.

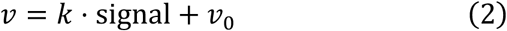

In our case, momentary motion or disparity evidence is integrated over time until it reaches one of two equidistant thresholds. The distance between these thresholds is governed by the boundary separation *a*. The starting point *z* determines whether the decision variable starts off closer to one threshold or the other. The influence of the perceptual evidence depends on the signed signal strength and a drift constant *k* according to **Equation 3**.

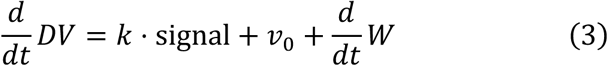

A Monte Carlo Markov Chain (MCMC) was run for 5000 iterations for each model with the first 200 iterations being discarded. The posterior probability distributions therefore reflect the 4800 iterations. Subsequently, 4 more repetitions were run to assess convergence of the chains, both qualitatively as well as using the formal Gelman-Rubin statistic (Gelman & Rubin, 1992). This lay between 0.99 and 1.01 in all cases except one (1.013), showing overall excellent convergence.

To simulate the data for a posterior predictive check, we used the built-in code within the HDDM package which simulates RT using discretized Wiener diffusion. We sampled 100 times from the MCMC trace of one model for each stimulus value and subsequently fed the parameters at those iterations into the function which estimated the first passage times of the Wiener process using an algorithm provided by (Navarro & Fuss, 2009). A second DDM run was performed on trials identified by the GLM-HMM with the highest posterior probability for each state where the number of trials was qualitatively deemed sufficient per difficulty (at least 50).

### Dimensionality reduction

A supervised dimensionality reduction method was utilised to identify the main differences in the estimated DDM parameter space between the experimental conditions. We used the estimated parameter distribution derived from the 5 runs of the MCMC for the human data as input data. LDA produces *C* − 1 dimensions for *C* classes which optimally discriminate between them by simultaneously minimising within-class variance and maximising between-class variance. Numerically, this comprises diagonalisation of the within-class and subsequent between-class covariance matrices. A Python-based implementation of this procedure is available using the *Scikit-learn* package (Pedregosa et al., 2011).

### GLM-HMM

We followed closely the procedure outlined in (Ashwood et al., 2022) using the open-source Python-based code. Briefly, we created a GLM with the logistic function as link function predicting rightward choices and including several covariates; we explored the previously used covariates bias, previous choice, stimulus strength and based on log-likelihood analysis included all three. We used 5-fold cross-validation for humans and *K*-fold cross-validation for monkeys, based on the *K* number of sessions for each animal, and derived the log-likelihood from averaging over the log-likelihood obtained in each hold-out set for all iterations for the different models with a varying number of states to determine the best one. The log-likelihood itself was used by the expectation-maximisation algorithm as a means to estimate the posterior probability of the GLM-HMM parameters given the data by virtue of maximum likelihood estimation (MLE). It refers to the likelihood of observing the specified dataset (choice) given the set of parameters. In order to understand this quantity more meaningfully, we divided this quantity by the number of trials in order to obtain the log-likelihood per trial. On the basis of steepest change in log-likelihood, we used three states for the global fits. The model itself is fitted with the expectation-maximisation algorithm. Different initializations converged on the same three states almost without exception. Only the first 1029 trials were included in the human data GLM-HMM analysis due to a shortage of trials for some participants. As a control for any imbalance in the saccade/hand trials, we fitted separately each of the four human data groups (cylinder/saccade; cylinder/hand; RDK/saccades; RDK/hand). We found that the fitted states were almost identical for each pair with the same stimulus/different motor responses and confirmed the main difference between the two stimuli to be judged (**Supplementary Figure 5-8**).

## Code availability

All code used in the preparation of this article can be found at github.com/Revanchist317/DDM_rangotis_et_al.

## Acknowledgments

We thank Sarah Kriener for producing the stimulus videos, Jochen Braun for statistical analysis consultation, Nela Cicmil and Maria Rüsseler for animal training and behavioural data, Bashir Ahmed and Jackson Smith for help with the headpost implantation. This work was funded by the DFG (Heisenberg Professorship Project-ID 406269671; grants with Project-IDs 406679869 and 425899996 to K.K.) and the Wellcome Trust (101092/Z/13/Z; KK and AJP). KK is a PI in the SFB1436. Funding was also from the Max Planck Society and the Humboldt Foundation (PD). PD is a member of the Machine Learning Cluster of Excellence, EXC number 2064/1 – Project number 39072764 and of the Else Kröner Medical Scientist Kolleg “ClinbrAIn: Artificial Intelligence for Clinical Brain Research”.

---

**Supplementary Figure 2-1.**
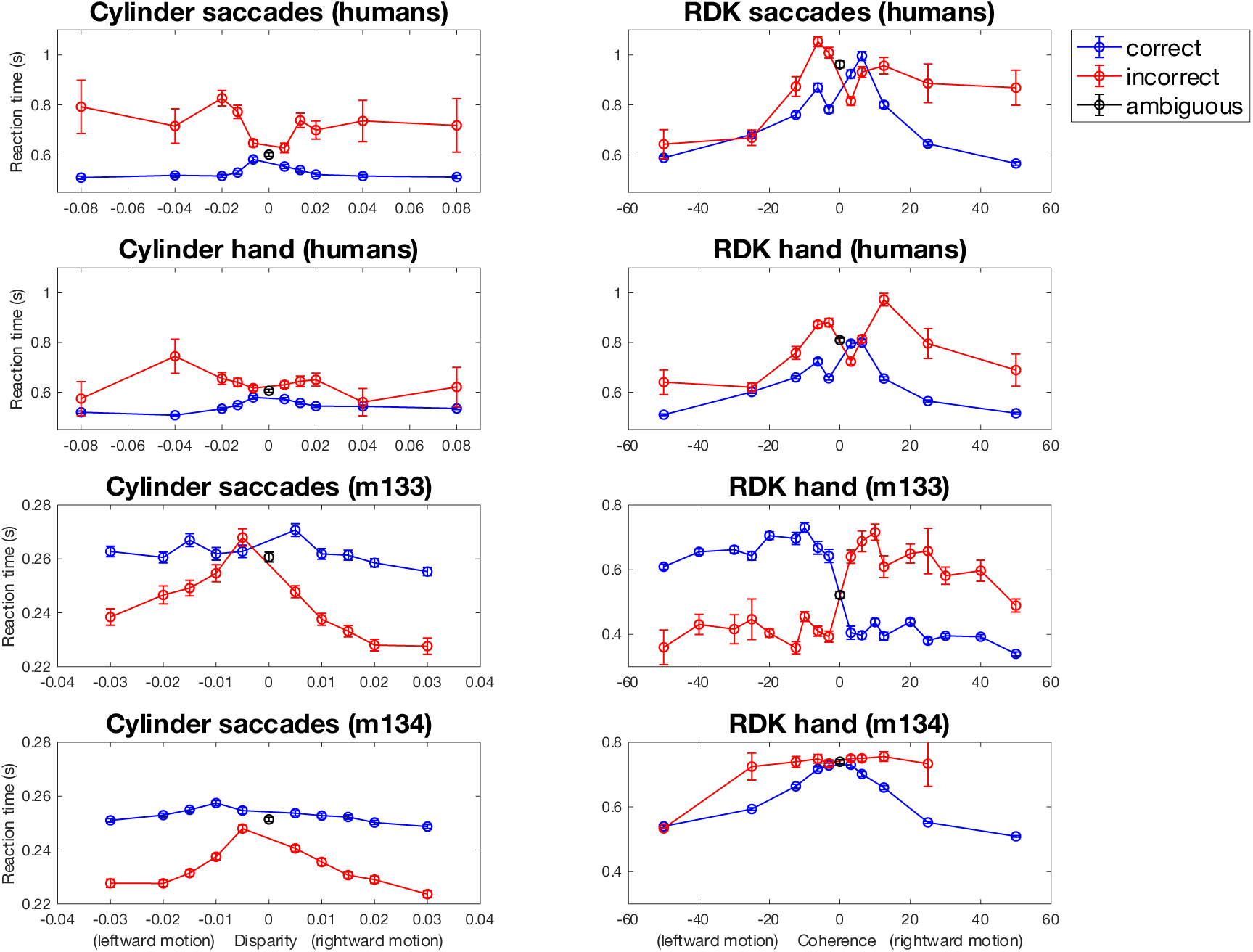
The distribution of reaction times (RT) in seconds against signal strength for each paradigm. The RT distributions show the expected pattern of slower responses for the more difficult stimuli around 0° (cylinder, left) and 0% (RDK, right). The RT distributions for correct choices are flatter for the cylinder tasks than in the RDK tasks for both species. Incorrect choices tended to be slower except for the saccade responses of the two monkeys. There is a clear bias shown by m133 for RDK with hand responses.

**Supplementary Figure 2-2.**
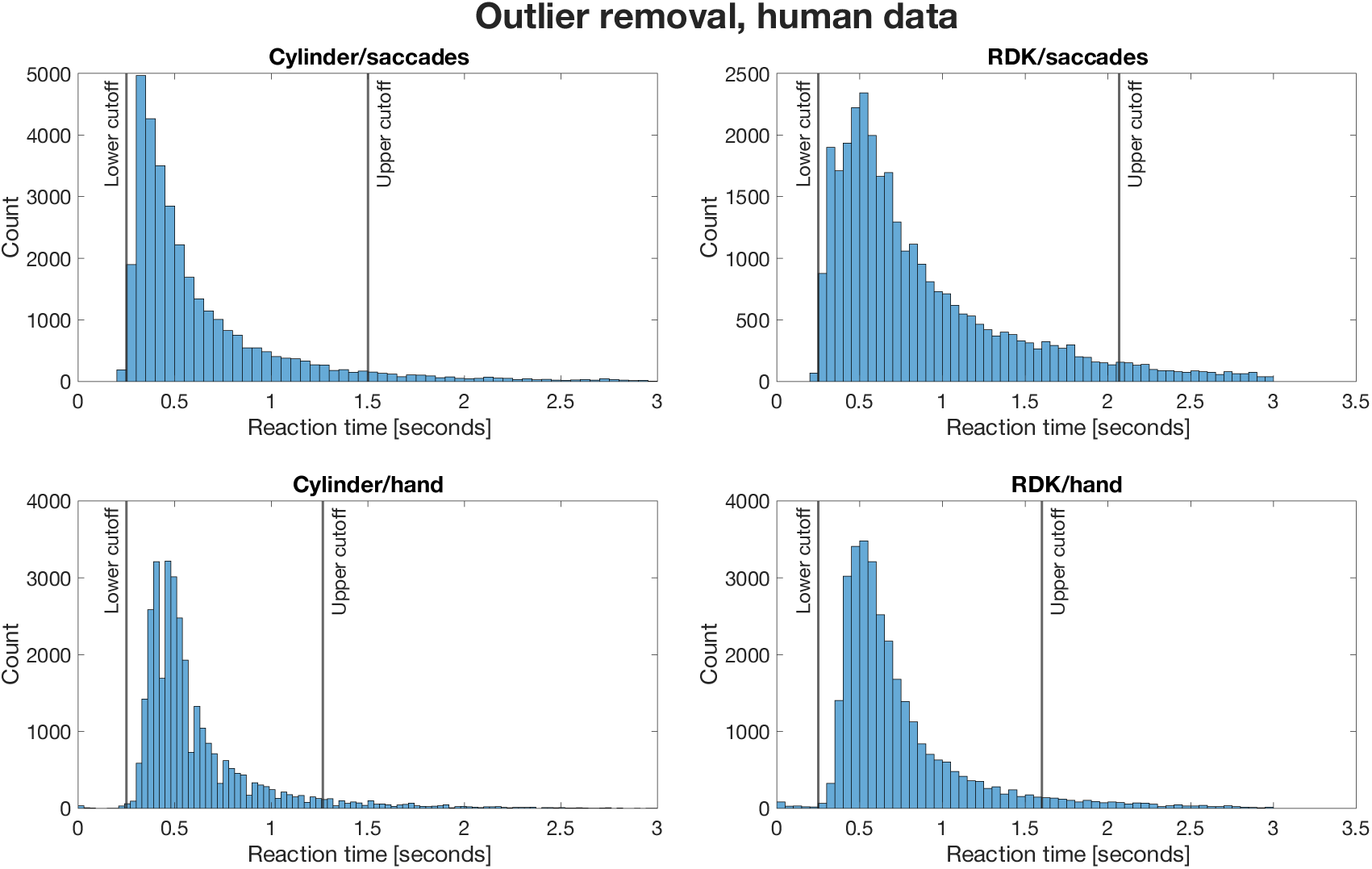
Outlier removal for the human RT data. Short reaction times below 250 ms were excluded as accidental responses or guesses and long reaction times (>95^th^ percentile) were excluded as they might be due to inattention. **Table 2** summarizes the cut off points for outlier removal for all reaction time data presented in the paper.

**Supplementary Figure 2-3.**
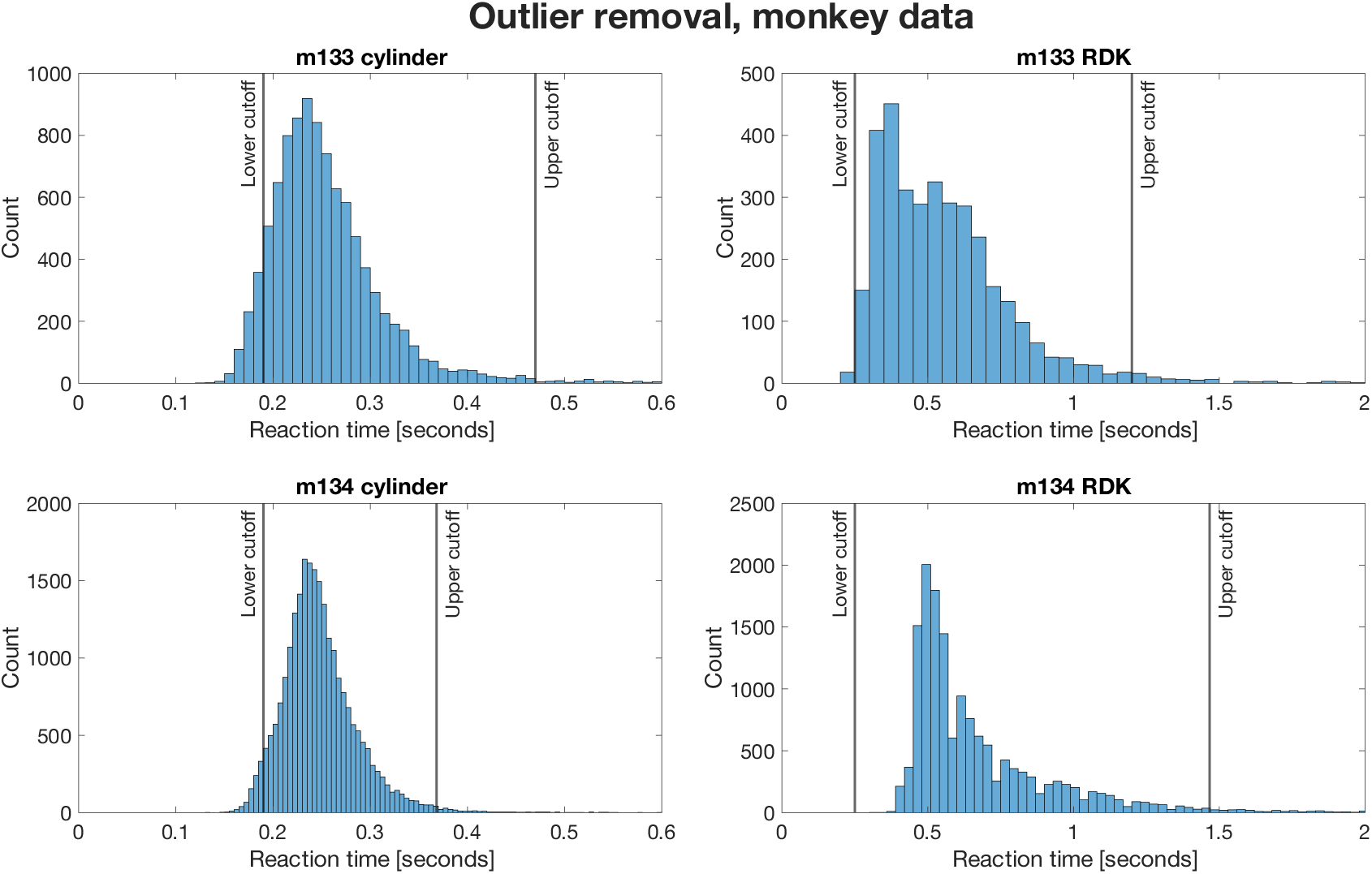
Outlier removal for the monkey RT data. Short reaction times (RDK: 250 ms, Cylinder: 190 ms) were excluded as accidental responses or guesses and long reaction times (> 98^th^-99^th^ percentile) were considered to be due to inattention and also excluded from the analysis (**Table 2**).

**Supplementary Figure 3-1.**
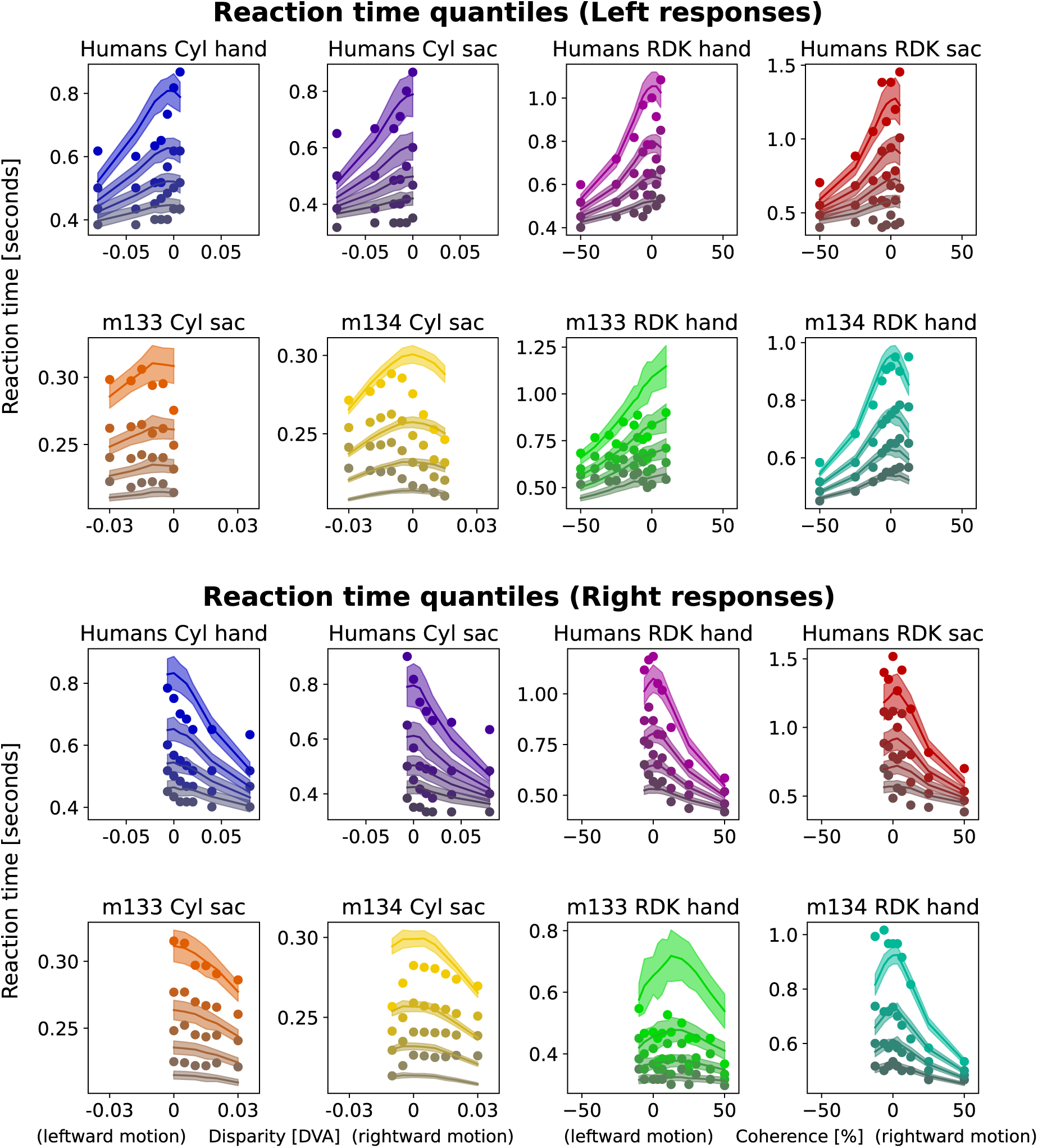
Qualitative goodness-of-fit comparison of the DDM simulations to the reaction time data. A posterior predictive check was performed to check model accuracy. We simulated artificial data using the derived parameters (solid lines and shaded regions to show +/- 1 SEM) and superimposed the RT quantiles from the empirical data (scattered dots). Conditions with less than 10 responses are not shown and were excluded from prediction. The model fit overall qualitatively well to the reaction time data based on a posterior predictive check but it could not fully account for the relatively long response times on the easiest trials in the cylinder task when compared to the ambiguous trials, consistent with the flatter RT pattern observed in **Supplementary Figure 2-1**.

**Supplementary Figure 5-1.**
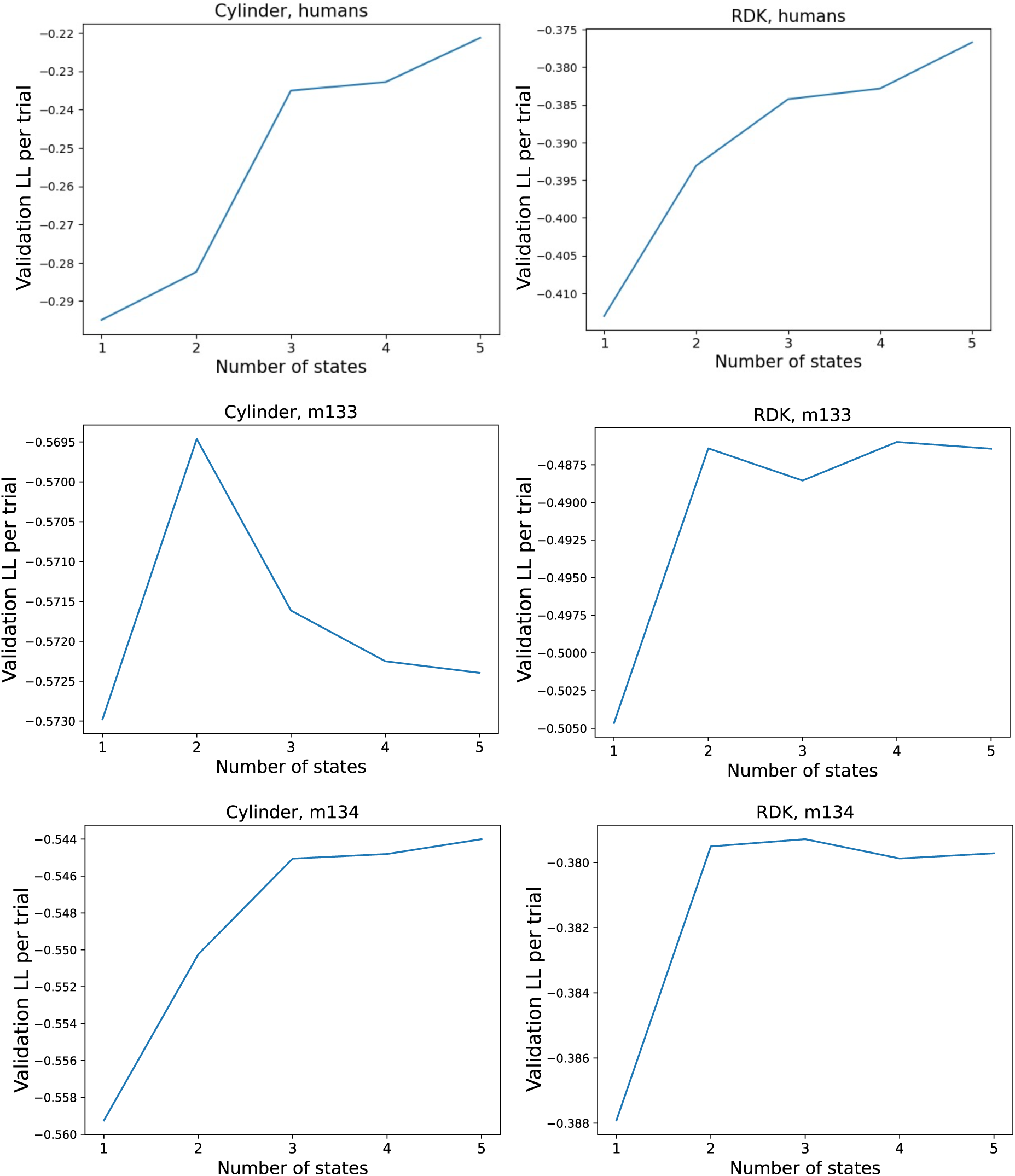
The validation log-likelihood per trial of observing the specific choices (left or right) given the set of parameters describing the total GLM-HMM is plotted as a function of the number of states for each GLM-HMM.

**Supplementary Figure 5-2.**
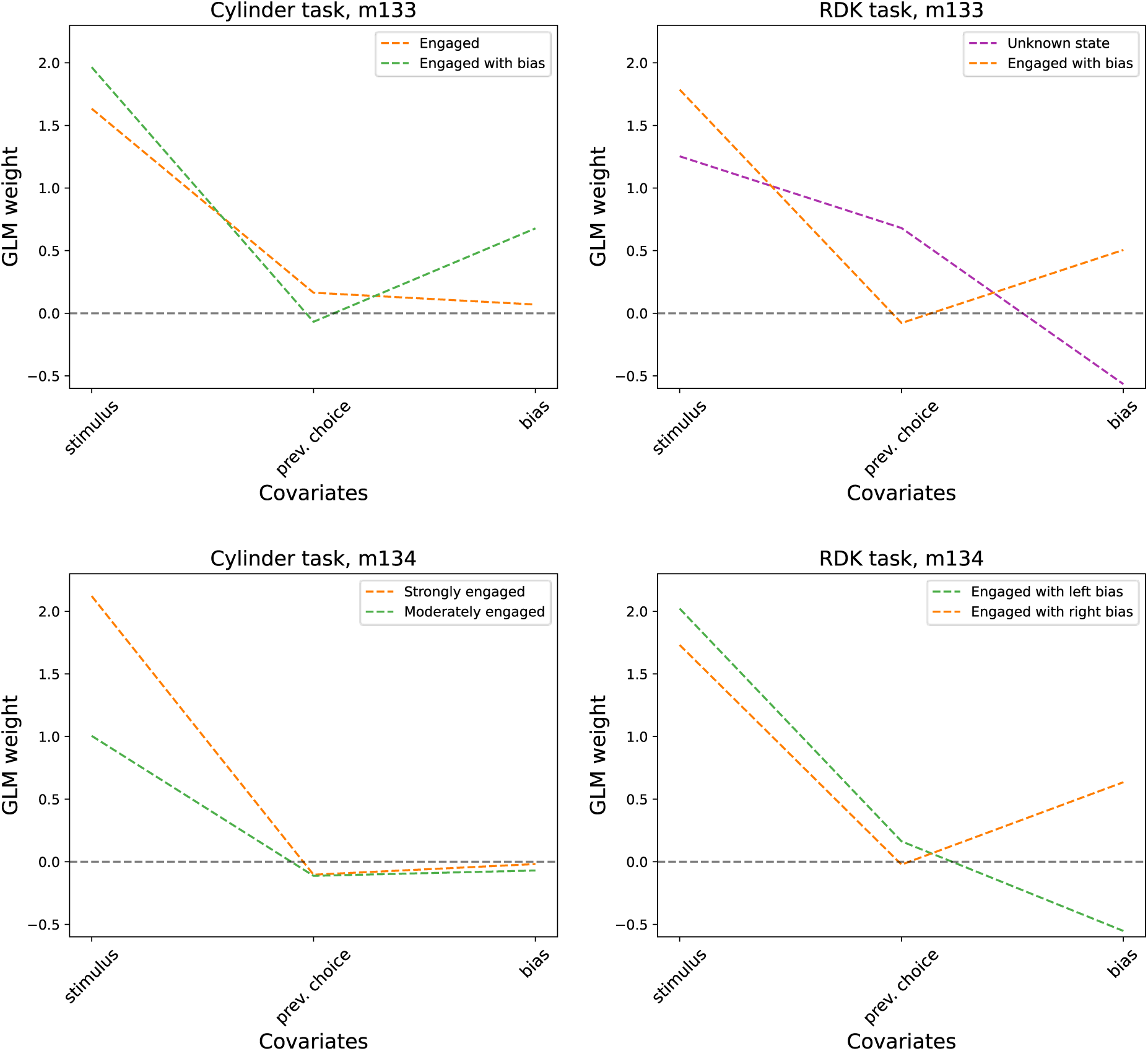
The results of the GLM-HMM analysis for the monkeys. The left panels show the states derived for the cylinder data and the right panels are the RDK data. Two states were derived based on the test log likelihood per trial result (see **Supplementary Figure 5-1**). The m134 GLM weights closely mirror those found for humans.

**Supplementary Figure 5-3.**
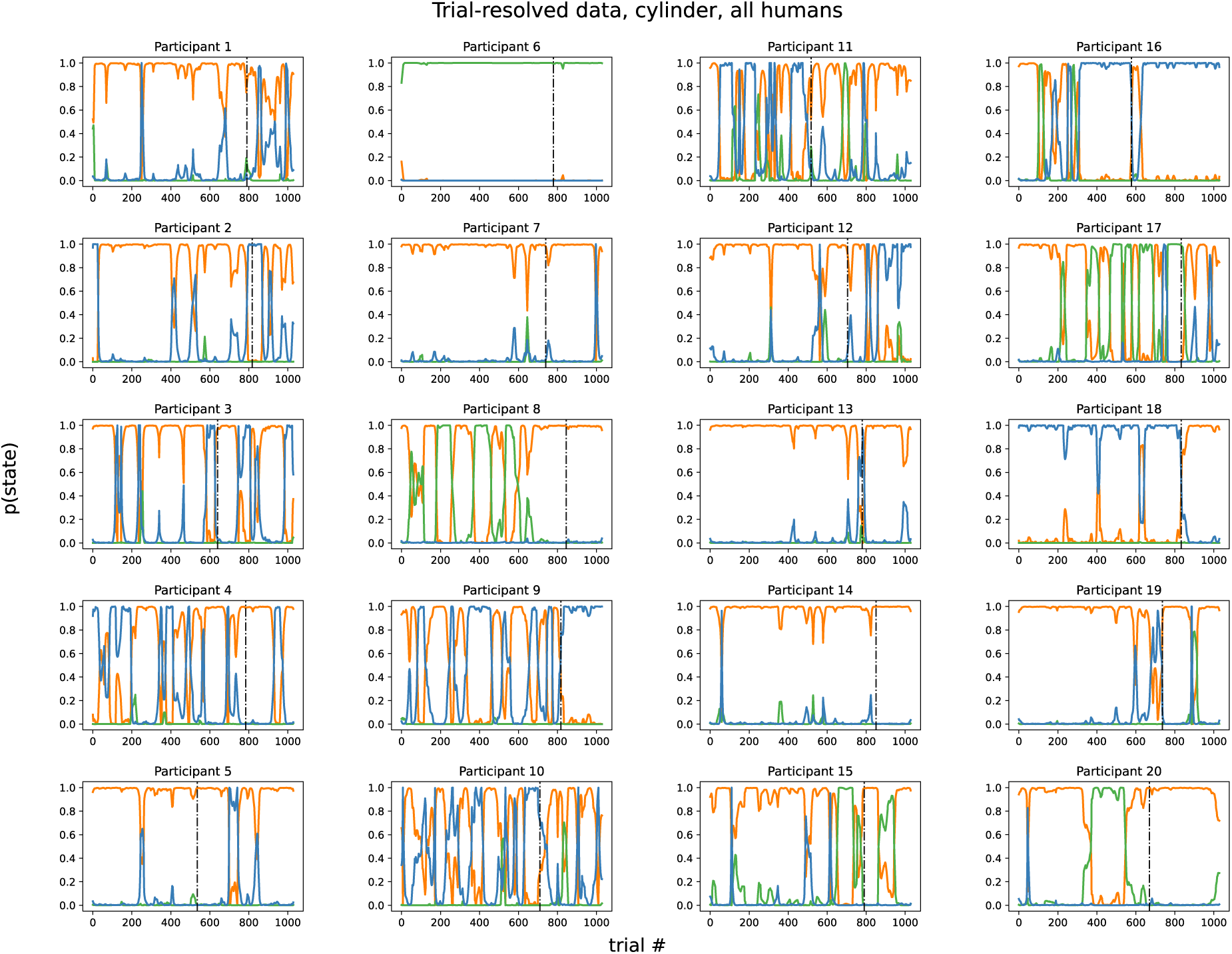
Trial-resolved GLM-HMM posterior probability data for the human participants on the cylinder task. The probability for specific states is shown trial-by-trial for each participant. The dash-dotted line separates trials with the hand responses on the left side and the saccadic response on the right side (see Methods). Traces are colour-coded by identified state in Figure 5A. (orange - strongly engaged; blue - moderately engaged with bias; green - disengaged).

**Supplementary figure 5-4.**
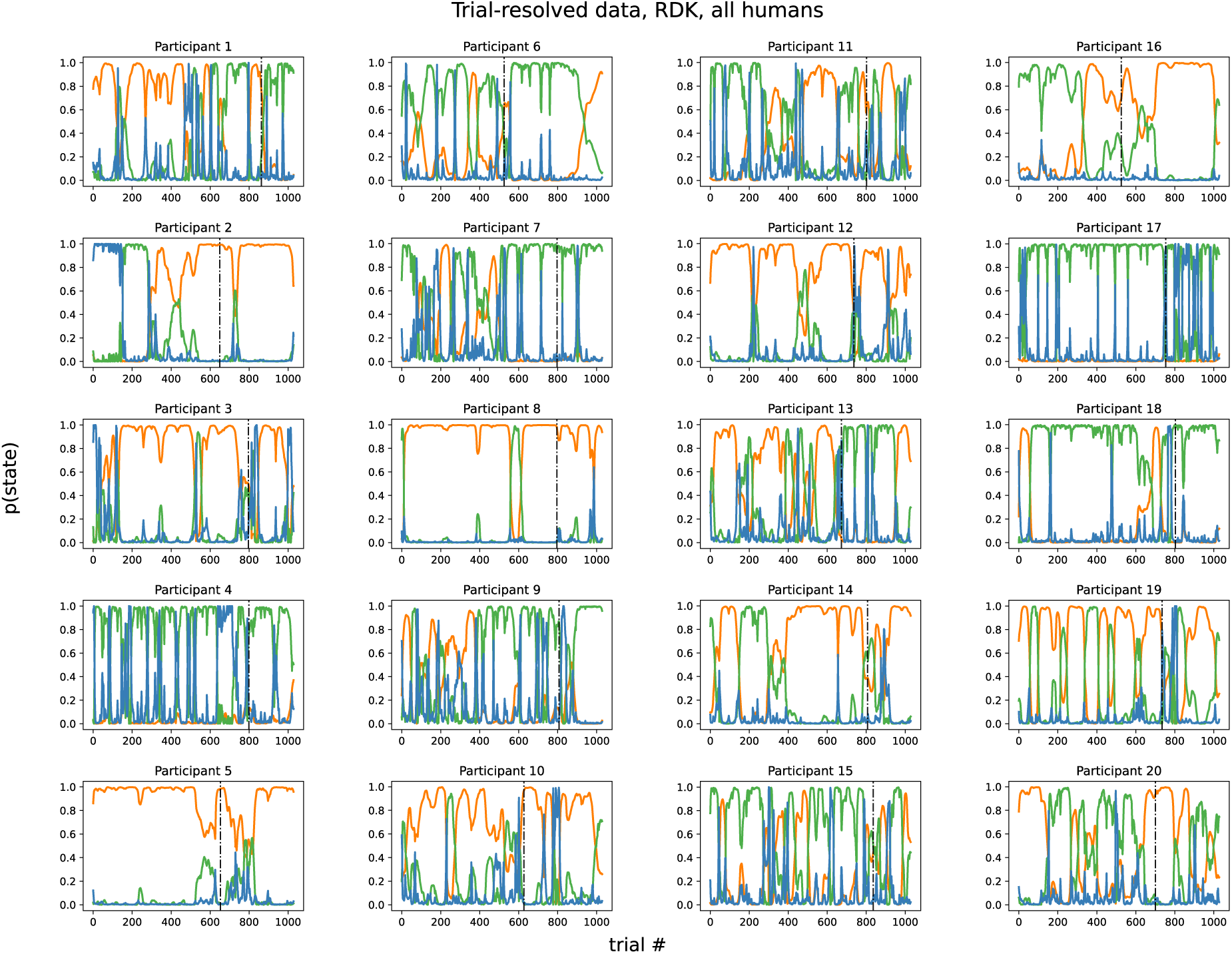
Trial-resolved GLM-HMM posterior probability data for the human participants on the RDK task. The probability for specific states is shown trial-by-trial for each participant. The dash-dotted line separates trials with the hand responses on the left side and the saccadic response on the right side (see Methods). Traces are colour-coded by identified state in Figure 5A. (orange - engaged; blue - engaged with left bias; green - disengaged).

**Supplementary Figure 5-5.**
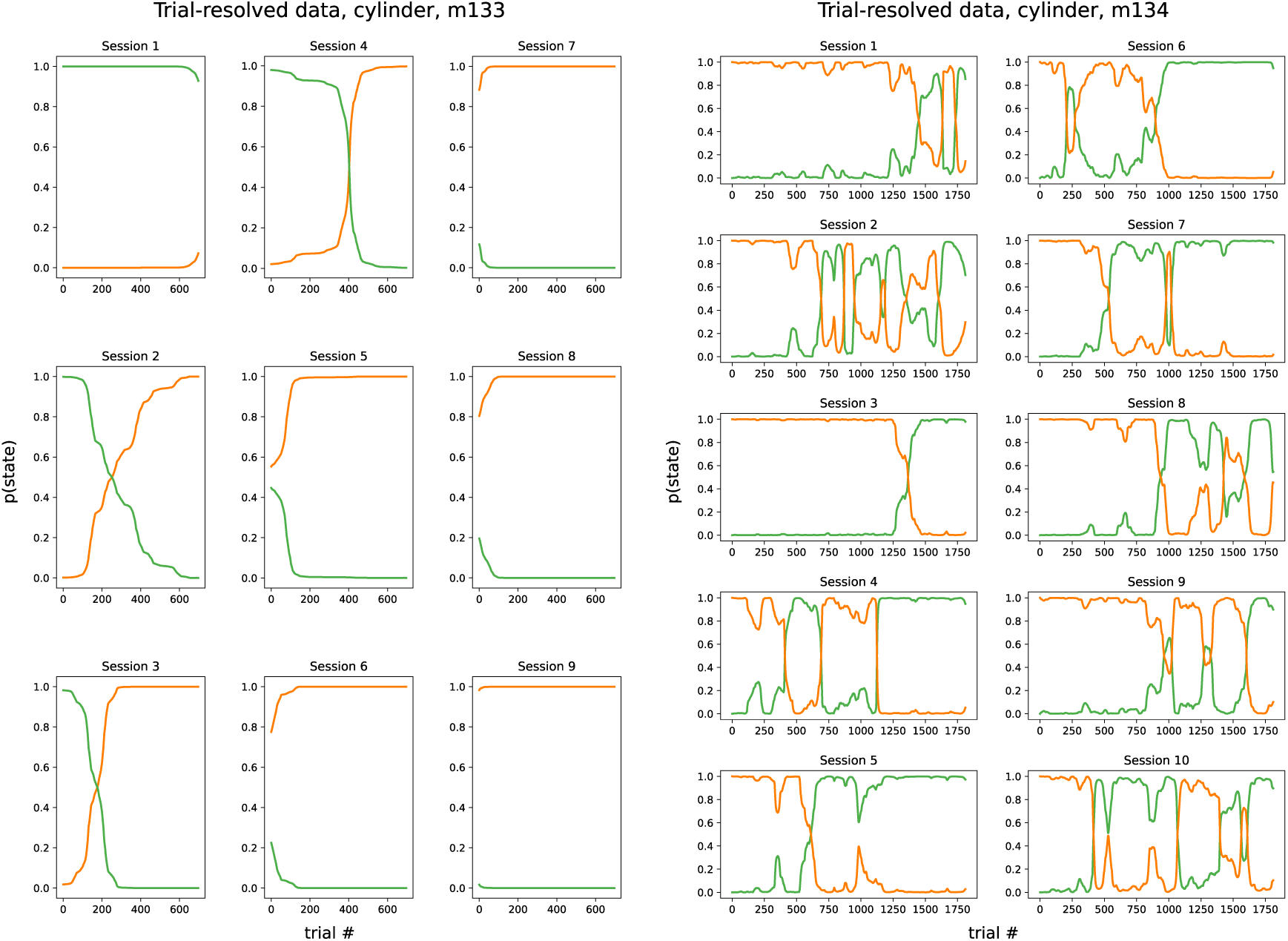
Trial-resolved GLM-HMM posterior probability data for the macaques on the cylinder task. The colours match the states identified in **Supplementary Figure 5-2.** While the pattern of trial-resolved data of m134 is switching similarly to many of human participants, m133 shows very few state-switches like only a couple of human participants for the cylinder task (see **Supplementary Figure 5-3).**

**Supplementary Figure 5-6.**
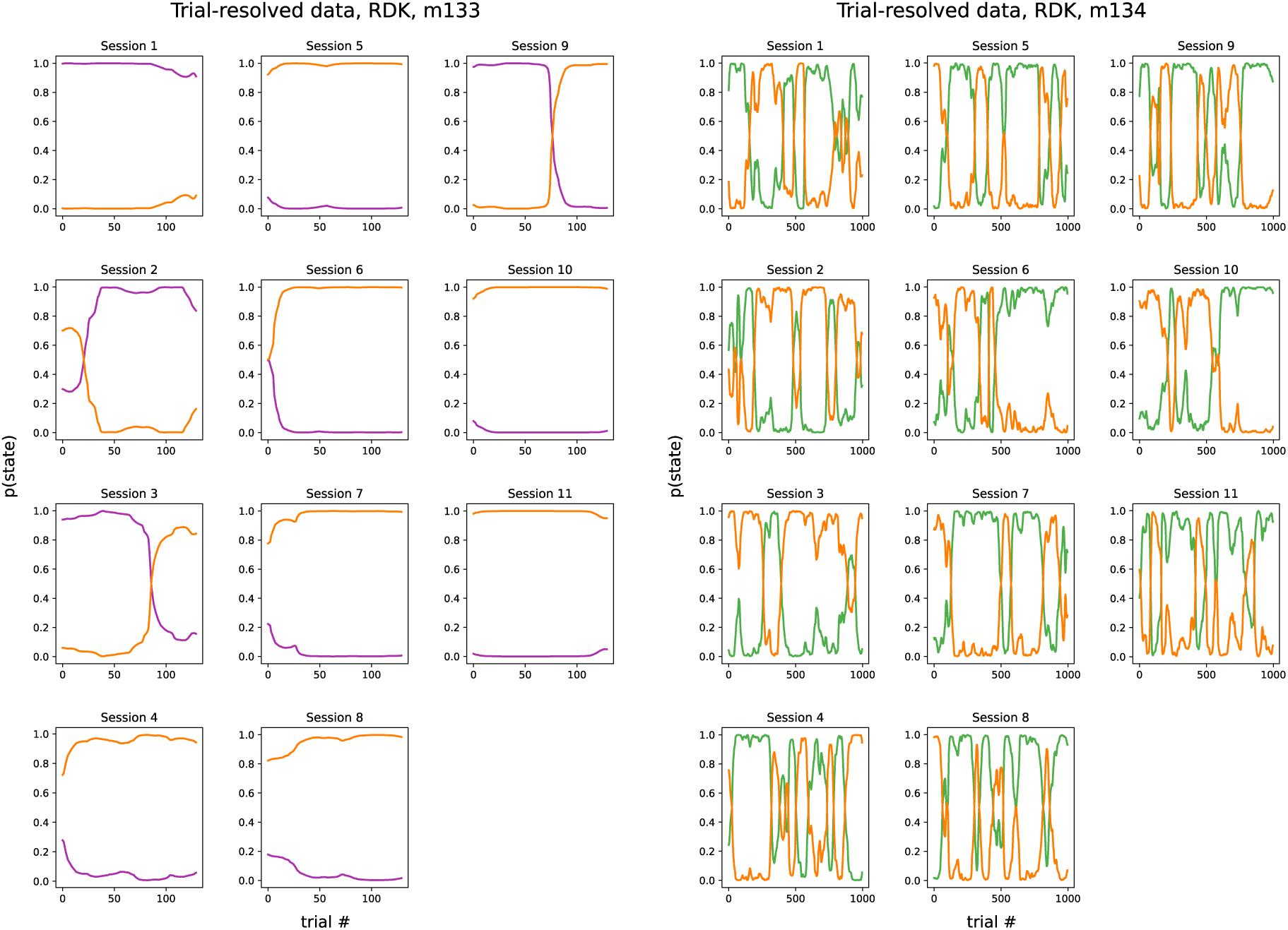
Trial-resolved GLM-HMM posterior probability data for the macaques on the RDK task. The colours match the states identified in **Supplementary Figure 5-2.** While the pattern of trial-resolved data of m134 is switching similarly to most of human participants, m133 shows very few state-switches like hardly any of human participants (see **Supplementary Figure 5-4).**

**Supplementary Figure 5-7.**
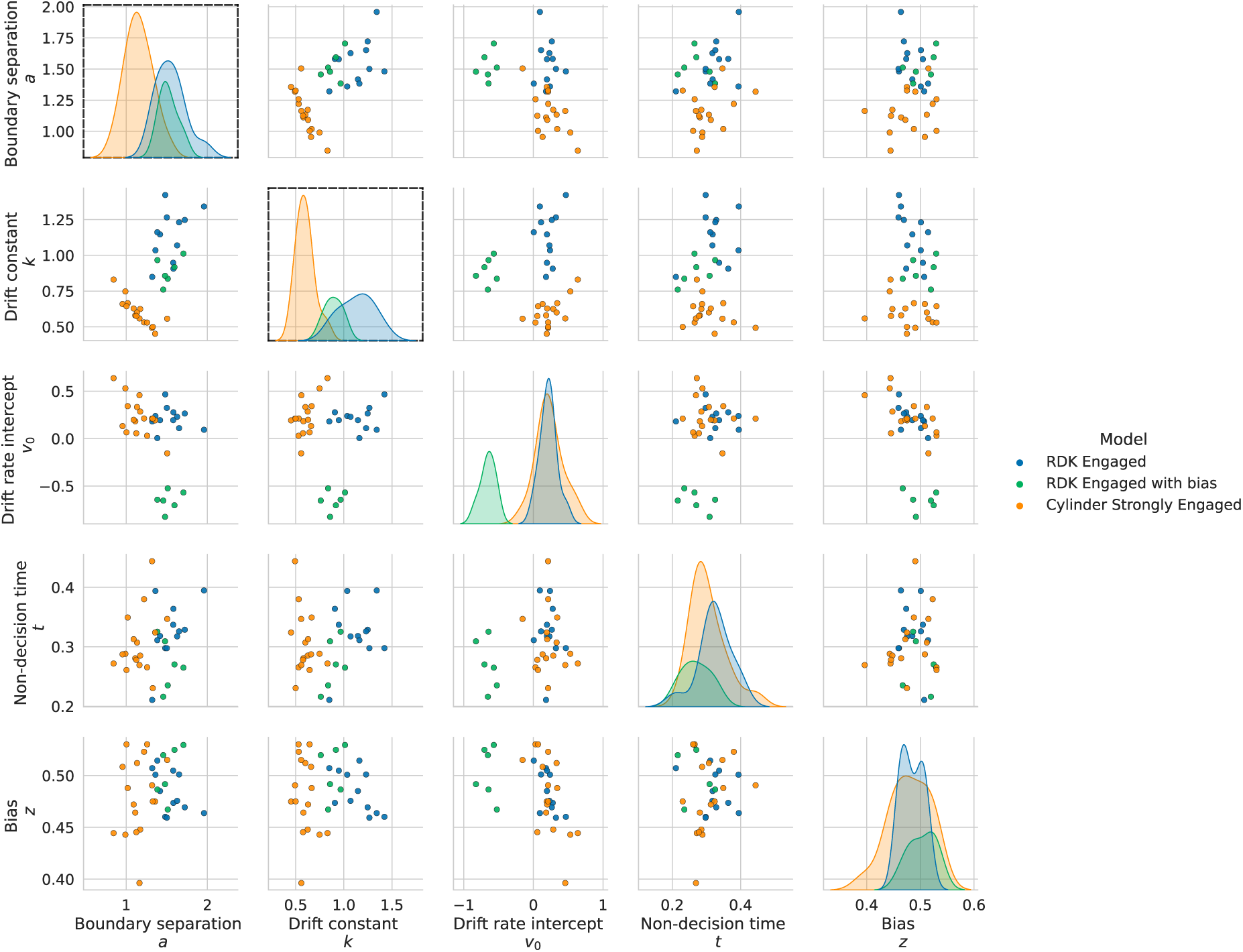
DDM of the trials from the human data set identified by the three main “engaged” GLM-HMM brain states. The DDM parameters show again the separation by stimulus type with a lower drift constant *k* and a lower boundary separation a for the cylinder stimulus versus the RDK for the three “engaged” states (black boxes). The bias identified in the green RDK state (“engaged with bias”) might be attributed to the difference in drift rate intercept (*v*_0_).

**Supplementary Figure 5-8.**
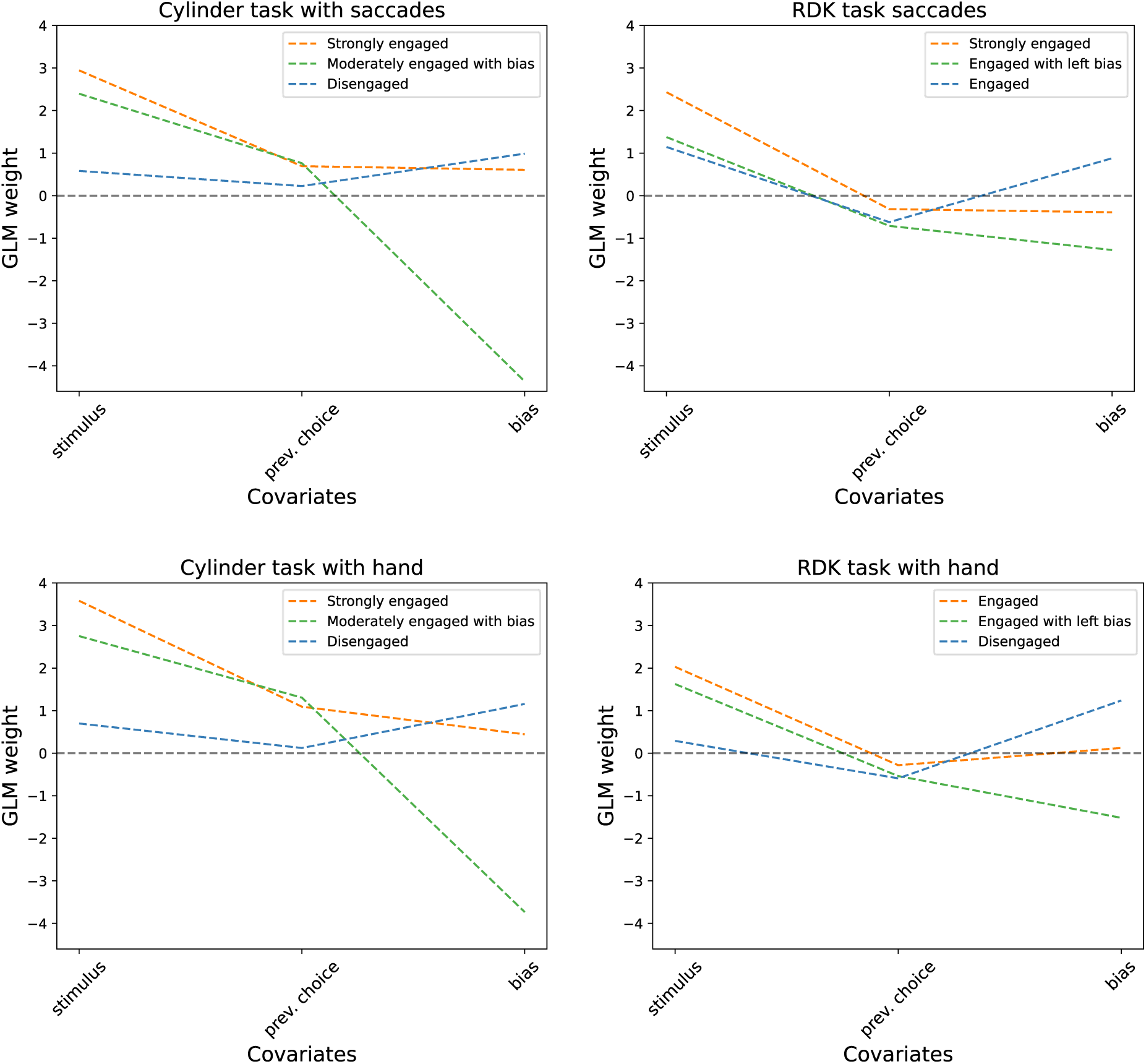
The GLM-HMM analysis for trials of human data grouped by the combinations of motor response and visual stimulus presented. The GLM weights suggest that the motoric response accounts for very little change between the estimated states.

